# Tracing the evolution of human gene regulation and its association with shifts in environment

**DOI:** 10.1101/2021.07.05.451164

**Authors:** Laura L. Colbran, Maya R. Johnson, Iain Mathieson, John A. Capra

## Abstract

As humans spread throughout the world, they adapted to variation in many environmental factors, including climate, diet, and pathogens. Because many of these adaptations were likely mediated by multiple non-coding variants with small effects on gene regulation, it has been difficult to link genomic signals of selection to specific genes, and to describe the regulatory response to selection. To overcome this challenge, we adapted PrediXcan, a machine learning method for imputing gene regulation from genotype data, to analyze low-coverage ancient human DNA (aDNA). First, we used simulated genomes to benchmark strategies for adapting gene regulatory prediction to increase robustness to incomplete aDNA data. Applying the resulting models to 490 ancient Eurasians, we found that genes with the strongest divergent regulation among ancient populations with hunter-gatherer, pastoralist, and agricultural lifestyles are enriched for metabolic and immune functions. Next, we explored the contribution of divergent gene regulation to two traits with strong evidence of recent adaptation: dietary metabolism and skin pigmentation. We found enrichment for divergent regulation among genes previously proposed to be involved in diet-related local adaptation, and in many cases, the predicted effects on regulation provide explanations for previously observed signals of selection, e.g., at *FADS1*, *GPX1*, and *LEPR*. For skin pigmentation, we applied new models trained in melanocytes to a time series of 2999 ancient Europeans spanning ~38,000 years BP. In contrast to diet, skin pigmentation genes show little regulatory change over time, suggesting that adaptation mainly involved large-effect coding variants. This work demonstrates how aDNA can be combined with present-day genomes to shed light on the biological differences among ancient populations, the role of gene regulation in adaptation, and the relationship between ancient genetic diversity and the present-day distribution of complex traits.

## Introduction

In the last decade, the number of ancient DNA (aDNA) samples from anatomically modern humans (AMHs) has increased dramatically (Marciniak and Perry, 2017). These samples span the globe, and cover time periods from several hundred to tens of thousands of years ago. This is a rich data source for understanding genetic changes and adaptations that occurred as humans expanded across the globe. However, linking genetic differences in aDNA samples to phenotypes poses several challenges (Irving-Pease *et al.*, 2021). First, while the samples are often paired with archaeological information, this is limited to what biological material has survived for thousands of years. Thus, most phenotypes of interest are not directly measurable. Second, due the complexity of many phenotypes and gaps in our knowledge of the genetic architecture of most traits, drawing conclusions about most phenotypes of interest based on genetic information alone is challenging (Benton *et al.*, 2021; Li *et al.*, 2020).

To date, most studies have focused on comparing aDNA from different geographical regions to map migrations and their relationship to archaeological changes (Skoglund and Mathieson, 2018). Shifts from a hunter-gatherer lifestyle to pastoral herding and agricultural farming have been of particular interest, because these changes had profound implications for multiple aspects of life. These include changes in day-to-day activities, population density, interactions with the environment, and substantial dietary shifts, such as increased reliance on domesticated grains (Goude and Fontugne, 2016; Olsson and Paik, 2016). These shifts likely modified selective pressures on populations as their lifestyles, diets, and pathogen exposures changed.

Genomic scans in present-day populations have identified many loci with evidence of positive selection (Field *et al.*, 2016; Grossman *et al.*, 2013; Rees *et al.*, 2020; Voight *et al.*, 2006). In some cases, selection can be linked to changes in the coding sequence of specific genes (Grossman *et al.*, 2013; Lamason *et al.*, 2005). In others, it can be linked to changes in gene regulation. For example, selection at the *FADS1* locus is linked to increased expression (Buckley *et al.*, 2017; Mathieson and Mathieson, 2018; Ye *et al.*, 2017). However in most cases, the molecular basis of signals of selection remains poorly understood, even when a specific gene can be implicated. For example, the leptin receptor (*LEPR*) is surrounded by a haplotype that has experienced recent positive selection (Voight *et al.*, 2006), and protein-coding changes in *LEPR* have been implicated in increased cold tolerance (Hancock *et al.*, 2008). However, altered expression of this gene is also associated with altered appetite regulation and metabolism (Kentish *et al.*, 2013; Loos *et al.*, 2006). Due to the difficulty in measuring environmental variables and disentangling LD patterns, it remains unclear whether selection is acting on coding variants, expression changes, or both, and which environmental variable is the source of the selective pressure (Luca *et al.*, 2010). Even these examples are exceptional; most selection signals cannot even be confidently attributed to specific genes. Selection peaks often span many genes, with little indication of which might drive changes in fitness or the underlying molecular mechanisms. This motivated us to ask whether information about variants associated with gene expression, such as expression quantitative trait loci (eQTL), could help to identify genes under selection—analogous to the way in which eQTL data can inform variant-gene-phenotype mapping in genome-wide and transcriptome-wide association studies.

We therefore developed an approach to identify genes whose regulation shifted in coordination with lifestyle changes in recent human history. These differences in regulation between ancient human groups in distinct environments suggest adaptation. To quantify gene regulation from aDNA samples, we adapted the PrediXcan-based approach we previously used to study gene regulation in archaic hominins (Colbran *et al.*, 2019; Gamazon *et al.*, 2015). Since available human aDNA have variable quality and coverage, we conducted simulations and control analyses to evaluate how models for imputing gene regulation perform when applied to low-coverage data, and how to ameliorate the effects of missing variants. These yielded heuristics for determining when regulation could be accurately modeled.

Guided by these simulations, we applied PrediXcan models for thousands of genes to hundreds of ancient humans representing populations from hunter-gatherer, pastoral, and agricultural lifestyles. We found enrichment for metabolic and immune pathways among the genes most divergently regulated between lifestyle groups. This reflects both the altered metabolic requirements and immune pressures of lifestyle shifts and highlights specific genes and pathways involved. For example, divergent regulation of *LEPR* suggests that its functions in metabolism and appetite regulation were relevant for recent adaptation. We also analyzed the predicted regulation of 20 diet-related genes in genomic regions with evidence of recent local adaptation. Supporting the accuracy of our approach, we rediscover the *FADS* locus regulatory haplotype that has been previously shown to vary by lifestyle and is likely the target of selection. We also identified divergent regulation between aDNA samples for selected genes involved in response to selenium (*GPX1*) and carnitine (*SLC22A5*) levels.

Modeling gene regulation using aDNA also allows us to characterise the nature of selection on specific phenotypes. To illustrate this, we investigated changes in predicted regulation of genes involved in skin pigmentation—the phenotype that is most clearly under directional selection in these populations—using PrediXcan models trained on expression data from melanocytes. We find that skin pigmentation genes show no consistent change in regulation over time suggesting that, for this particular phenotype, evolutionary change was driven by coding variants rather than regulatory changes. Overall, this work provides an atlas of imputed regulation for hundreds of ancient humans across thousands of genes to facilitate future exploration of gene regulatory shifts in recent human evolution, and demonstrates the utility of combining molecular predictive models with ancient DNA to understand the evolution of complex traits.

## Results

### Gene regulatory patterns can be imputed using low-coverage aDNA data

The genetically regulated component of gene expression can be predicted by machine learning models trained on gene expression. Previous approaches have applied these models to genome-wide common variant data from present-day humans (Fig. 1A), for example to perform transcriptome-wide association studies (Gamazon *et al.*, 2015; Zhou *et al.*, 2020; Zhu and Zhou, 2020), and to high-coverage archaic hominin genomes (Colbran *et al.*, 2019). Here, we adapt this approach to enable application to low-coverage genotype data from ancient human individuals, considering the unique attributes of these data. In particular, aDNA data vary in coverage, depth, and quality. This creates a trade-off between number of individuals available for analysis and the genotype quality.

**Figure 1:**
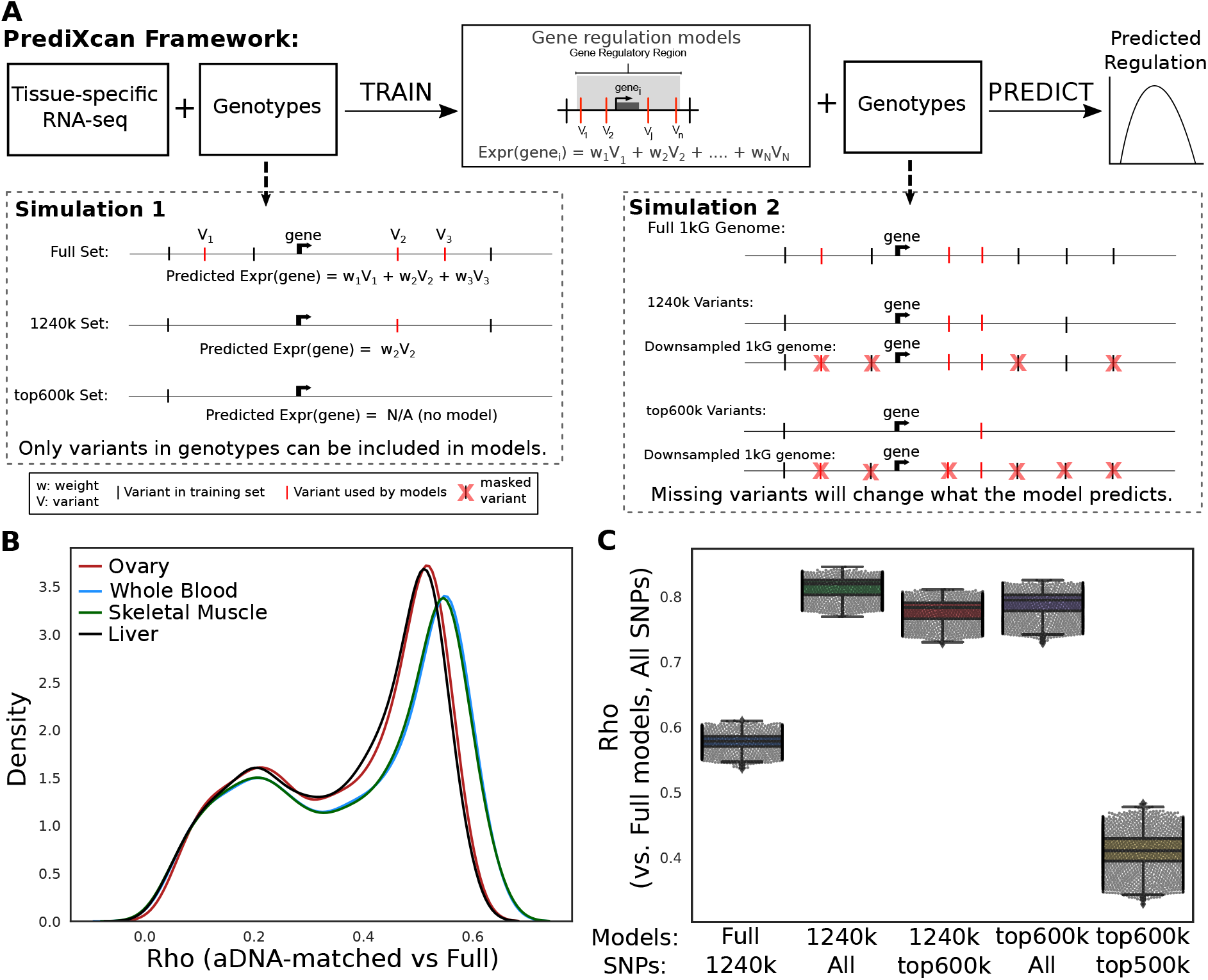
Gene regulatory prediction models can be trained for application to low-coverage ancient DNA. (a) Schematic of the framework for training and testing PrediXcan models. PrediXcan consists of statistical models for imputing genetic regulation of gene expression that are trained on genetic variants and normalized transcriptomes from diverse tissues collected as part of the GTEx Project. For each gene, PrediXcan considers genetic variants within 1 Mb of the gene (grey box) and uses elastic net regression to learn a combination of variants and weights to predict variance in its expression across individuals. Variants included in the final model are illustrated by red vertical lines. (b) To evaluate the potential for gene regulatory prediction using aDNA, we performed several analyses. First, we evaluated the effects of using three different variants for model training: Full (all common variants in GTEx), 1240k (all variants in the aDNA 1240k capture set), and top600k (the 600k most representative variants from the 1240k capture set; see Methods). We also simulated the presence of missing data in the prediction phase by masking variants from genomes from the 1000 Genomes project such that only variants from each of the 3 sets (Full, 1240k, top600k) were available for use in prediction. (c) Distribution of Spearman *ρ* between predictions per individual in four tissues (Skeletal Muscle, Whole Blood, Liver, Ovary) when considering the complete genome vs. 1240k-matched simulated ancient genomes. (d) Spearman *ρ* between predictions from a range of targeted models on down-sampled genomes to the Full PrediXcan models applied to all variants available for 1kG individuals. Models were trained on different variant subsets (x-axis, top row: All, 1240k, top600k) and applied to complete or downsampled 1kG genomes (x-axis, bottom row: All, 1240k, top600k). There is one point per individual sample.

To explore this trade-off and the feasibility of this approach on available aDNA data, we created simulated ancient genomes by removing variants from present-day individuals with whole-genome sequencing from the 1000 Genomes Project (1kG) (The 1000 Genomes Project Consortium, 2015). (See the Supplementary Materials for detailed discussion.) First, we found that PrediXcan models trained using common variants identified from present-day whole genome sequencing data are robust to random patterns of missing data (Spearman *ρ* > 0.75 with up to 45% of variants missing; Supplementary Fig. 2A). However, nearly all aDNA samples used here were genotyped by targeted capture of ~1240k variants (“1240k set”; Supplementary Fig. 1B) (Fu *et al.*, 2015; Haak *et al.*, 2015). Furthermore, many of the ancient samples have low genotyping coverage resulting in many missing variants (Supplementary Fig. 2B). Thus, we next matched the missing data to patterns observed in aDNA and compared the performance of different prediction models applied to full genomes vs. genomes with simulated missing data (Supplementary Fig. 1A). These models’ consistency decreased substantially when applied to genomes with missing data matched to that in ancient DNA samples (median Spearman *ρ* = 0.39; Fig. 1B).

To address this, we trained prediction models using only variants from the 1240k set. The predictions of these models were correlated with those of the full models (median Spearman *ρ* = 0.67), as expected given the LD between variants in the 1240k set and those in the full models. We also identified a set of variants that were most frequently available in the highest quality ancient samples; this resulted in a set of the 600,000 most-informative variants from the 1240k set (“top600k set”). We then trained models using these variants targeted to the aDNA data (top600k) and evaluated their performance on full genomes and simulated ancient genomes (Methods). While predictions made by the 1240k and top600k models were largely consistent with those made by the Full models when applied to genomes with no missing data (median rho 0.82 and 0.79 respectively), only the 1240k models maintained consistency when applied to incomplete genomes (Fig. 1C). We therefore concluded that the 1240k trained models strike a balance between accuracy and sample size when applied to ancient data, and thus we used these models for the rest of our analysis.

### Imputing gene regulatory differences between ancient human populations

We collected ancient human samples with genetic data from a variety of sequencing and genotyping platforms (Methods). Based on the analyses in the previous section, we ranked individuals by the number of sites successfully genotyped, and took the top quartile of individuals (>771240 SNPs, or 0.74x coverage), restricting to individuals from Eurasia due to sample density and genetic similarity to the training data (Fig. 2A). The samples ranged in date from 90 years before present (yBP) to 45,000 yBP, with the majority between 2,500 and 6,000 yBP (Fig. 2B).

**Figure 2:**
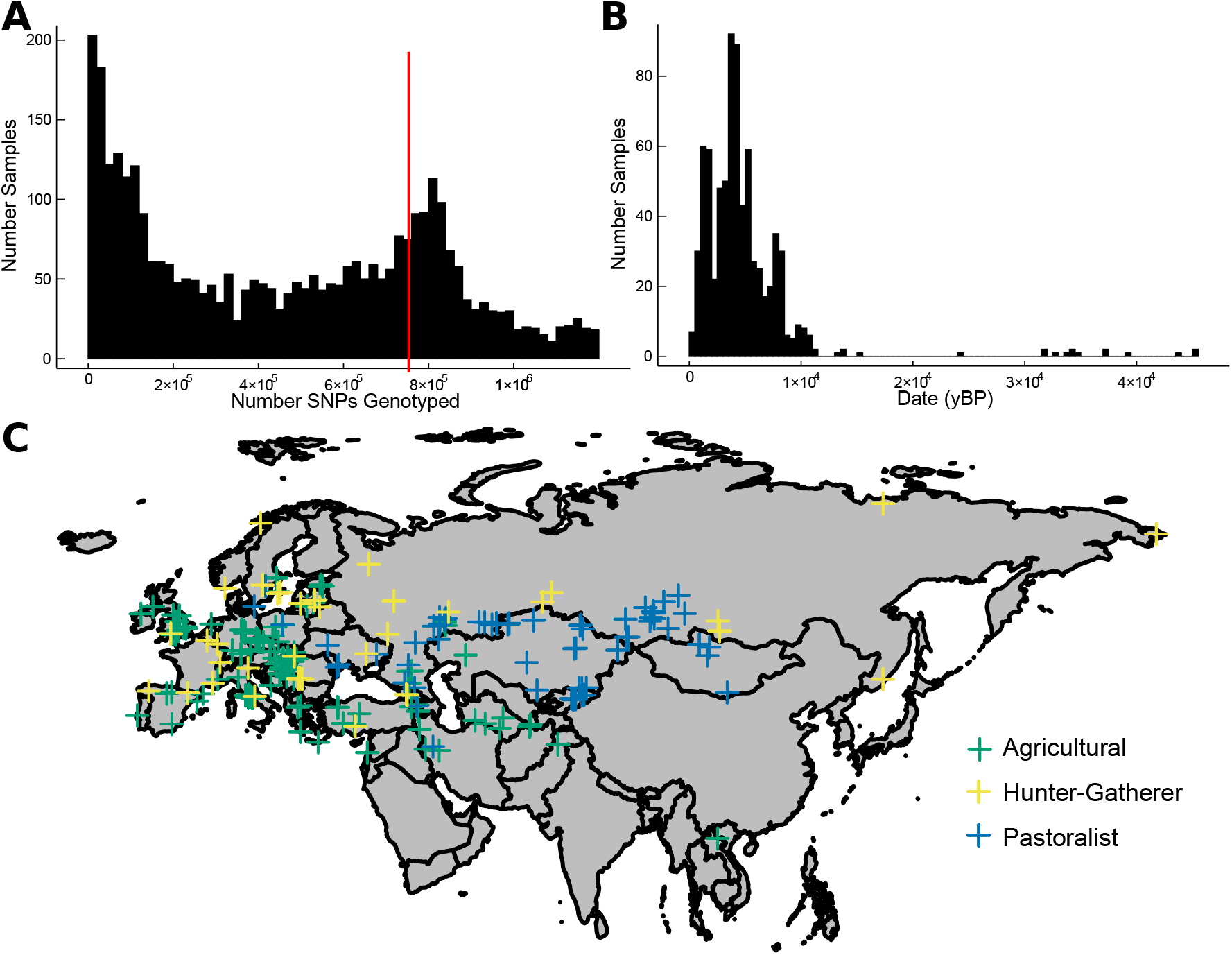
Attributes of ancient humans considered in this study. (a) Distribution of the number of variants with genotype call in the aDNA samples. The maximum is 1,233,013, the number of SNPs on the 1240k genotyping chip. We analyzed individuals in the 3rd quartile or above (red line, 771,029 SNPs). (b) Distribution of the age of 490 Eurasian samples analyzed in years before present (yBP). (c) We assigned ancient Eurasians with sufficient genetic data to three lifestyles: Green = agriculturalist, Blue = pastoralist, Yellow = hunter-gatherer.

We then assigned individuals to a lifestyle (hunter-gatherer, pastoralist, or agricultural) by literature review of the associated archaeological culture based on information from the original aDNA publications. In general, hunter-gatherers were from sites: 1) dated to times before any evidence of domestication or 2) with evidence only for foraging and meat consumption and no domesticated plants or animals. Agriculturalists were from sites with evidence for domesticated grains and animals. Pastoralists can be difficult to distinguish from agriculturists, and here refers to individuals from often semi-nomadic societies focused on domesticated animals (primarily the Yamnaya and similar groups). In addition, in some cases, the lifestyle distinction was based on genetic similarity to other groups, so the categories used here are based on a combination of genetics and archaeology. Because of these difficulties, we focused primarily on comparisons between hunter-gatherers and the other groups. This process resulted in 490 ancient Eurasian individuals with an assigned lifestyle and aDNA for further study (Fig. 2C).

We then applied the 210,800 “1240k” gene regulation prediction models described in the previous section to the 490 ancient samples, as well as to 503 present-day Europeans from the 1000 Genomes Project (The 1000 Genomes Project Consortium, 2015). This resulted in normalized expression predictions in different tissues (“predicted regulation”) for 14,873 unique genes. The observed expression level of a gene in an tissue in an individual is a combination of genetically regulated and environmental factors. The output of our prediction model is not a direct proxy for the observed expression, but rather a quantification of the genetic component of gene regulation. Thus, differences in predicted regulation between individuals reflect potential differences in the inherited genetic component of expression, not environmentally driven differences.

### Divergently regulated genes are enriched for immune and metabolic functions

To survey high-level differences among ancient individuals from the three lifestyle groups, we identified divergently regulated genes as those for which the *P*-value of a Kruskal-Wallis test passed a Bonferroni multiple testing correction (per-tissue). For example, *GPR84* was among the most differences in predicted regulation between populations (Fig. 3A; Adrenal Gland predicted regulation of −0.0421 in agriculturalists vs. 0.197 in hunter-gatherers; K-W *P* -value). Overall, 5759 unique genes showed evidence of divergent regulation between lifestyles in at least one tissue (median 2 tissues; Supplementary Fig. 6A), and an average of 9.8% of genes in each tissue were divergent (Supplementary Fig. 6B). However, most divergent genes had relatively small changes in magnitude between groups (e.g. maximum 1.17 magnitude difference between hunter-gatherers and agriculturalists in Subcutaneous Adipose) and the majority of these differences are likely attributable to genetic drift, rather than the effects of selection. We therefore imposed a genomic control (Methods) on the full distribution of 210,800 (genes × tissues) Kruskal-Wallis *P*-values (Fig. 3A). We focused on the 500 genes with the most evidence of divergent regulation (corrected *P* < 3.46 × 10^−3^, FDR = 0.586), which may be enriched for targets of selection.

**Figure 3:**
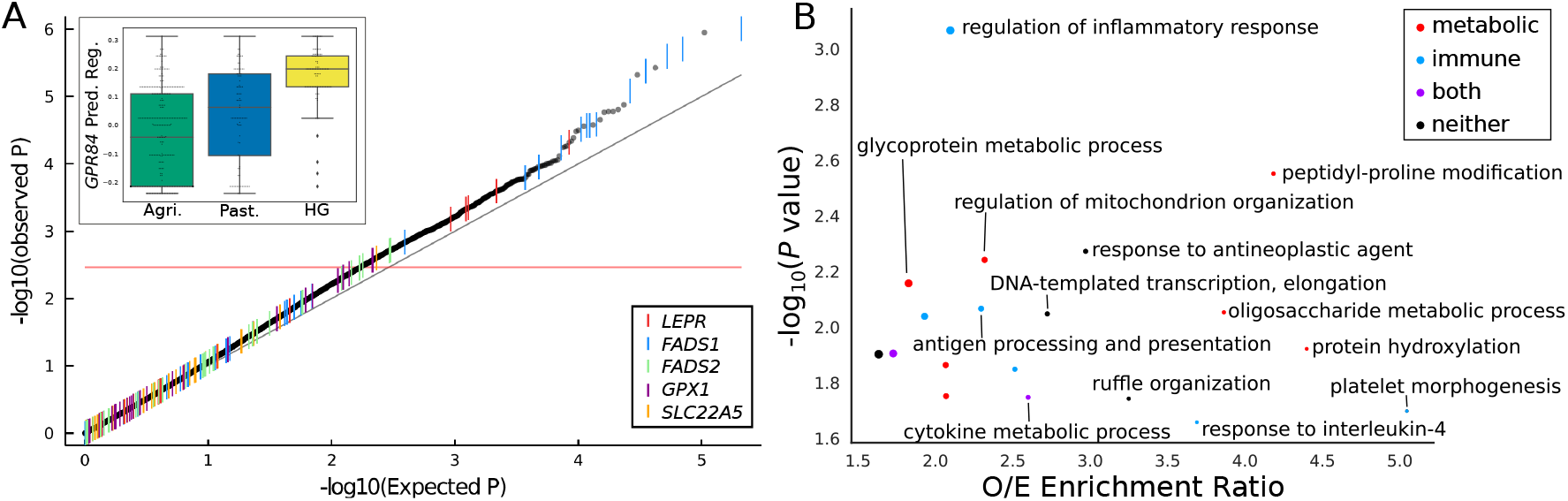
Immune and metabolic genes are among the most diverged between ancient lifestyle groups. (a) QQ plot for all gene regulation models in all tissues. Observed *P*-values are calculated after GC correction. The 500 most divergently regulated genes have at least 1 model above the red line. Inset: Predicted Regulation of *GPR84* in Adrenal Gland. (b) The most-enriched Gene Ontology (GO) terms among the 500 most diverged genes. Point size scales with number of diverged genes in each category (range 3-24).

We hypothesized that immune and metabolic traits were among those under the most selective pressure as populations transitioned between lifestyles. To identify systematic patterns in the 500 most divergently regulated genes, we conducted Gene Ontology (GO) over-representation analysis. The twenty most-enriched annotation terms (Fig. 3B) included immune-related (e.g.“antigen processing and presentation”) as well as basic metabolic processes and cellular functions (e.g. “glycoprotein metabolic process”). In addition, the enrichments for some of the more general terms may be driven by genes with pleiotropic immune system effects. For example, the eight genes driving the enrichment of the “DNA-templated transcription, elongation” term included *THOC5*, which also functions in immunity and response to stimuli through cytokine-mediated pathways (Mancini *et al.*, 2004; Tamura *et al.*, 1999), *ELP1,* which has functions in proinflammatory signalling (Cohen *et al.*, 1998), and *AFF4*, a component of the super elongation complex, which is recruited in response to HIV-1 infection (Chou *et al.*, 2013; He *et al.*, 2010).

Many gene sets are likely to maintain similar regulatory patterns across populations, regardless of lifestyle, and these should not be enriched among the top divergently regulated genes. To test this, we quantified the enrichment of three such sets under strong functional constraint among the 500 most diverged genes between lifestyle groups across tissues: 1) genes that have experienced stabilizing selection on their levels of expression across many species (Chen *et al.*, 2018), 2) genes responsible for core house-keeping functions (Eisenberg and Levanon, 2013), and 3) genes that are intolerant to loss-of-function coding variation (“LOF-intolerant”) in present-day humans (Lek *et al.*, 2016) (Methods). As expected, LOF-intolerant genes and those under long-term stabilizing selection are not enriched (Table 1). Surprisingly, housekeeping genes were slightly enriched (OR = 1.33, *P* = 0.0076). By definition, housekeeping genes have uniform and ubiquitous expression across tissues, so this pattern could partially be explained by increased power to model changes in their regulation in multiple tissues. However, many housekeeping genes are also involved in basic cellular metabolism (Eisenberg and Levanon, 2003), which could require fine tuning in response to changes in nutrient sources or other environmental shifts. We also tested for enrichment of genes that encode proteins that directly interact with viruses, since these genes are known to evolve rapidly (Enard *et al.*, 2016), but we find no enrichment among the top 500 genes, suggesting that selection at these loci could be driven by coding rather than regulatory changes.

**Table 1:** Skin pigmentation genes with nominally significant associations between ancient sample age and regulation. Betas and P-values were calculated using a linear regression of the predicted regulation on the date, including the first 10 ancestry principal components.

| Gene | $\beta$ (All) (95%CI) | $P$ (all) | $\beta$ (<15ky) (95%CI) | $P$ (<15ky) |
| --- | --- | --- | --- | --- |
| *TYR* | -2.05e-6 (-3.1e-6 - 9.67-7) | 0.00021 | -4.7e-6 (-6.43e-6 - -2.97e-6) | 1.08e-7 |
| *TRPM1* | -9.93e-7 (-1.97e-6 - -2.05e-8) | 0.045 | -2.039e-6 (-3.59e-6 - -4.83e-7) | 0.010 |
| *MITF* | 1.65e-6 (1.08e-7 - 3.2e-6) | 0.036 | 2.80e-6 (3.34e-7 - 5.27e-6) | 0.026 |
| *KIT* | -3.62e-7 (-7.56e-7 - 3.16e-8) | 0.071 | -7.08e-7 (-1.33e-6 - -8.10e-8) | 0.027 |

Several of the top divergently regulated genes underlying the GO functional enrichments have been implicated in local adaptation, for example *EP300* (Zheng *et al.*, 2017) and several subunits of HLA-DQ (Catassi and Catassi, 2018; De Silvestri *et al.*, 2018; Pierini and Lenz, 2018). In the next two sections, we explore the connection between sequence signatures of recent adaptive evolution and divergent gene regulation with a focus on diet and skin pigmentation.

### Changes in gene regulation contributed to adaptation to diet between ancient lifestyles

Many regions of the human genome bear signatures of recent population-specific adaptive evolution. However, the phenotypic drivers and molecular mechanisms underlying these evolutionary signatures are largely unresolved. Since diet was one of the main factors that shifted with the change from hunting and gathering to farming, we hypothesized that gene regulatory changes between lifestyle groups might be the target of signals of selection at dietary genes.

We compared the predicted regulation of 20 diet-related genes in regions with evidence of population-specific local adaptation (Rees *et al.*, 2020) between ancient human groups with different lifestyles (Methods). Models for the 20 genes tested were enriched for lower *P*-values (*P* = 1.19 × 10^−14^, K-S test), with 4 unique genes among the top 500 most diverged genes by group (Supplementary Table 4).

*FADS1* showed the most consistent evidence for divergent regulation between agriculturalists, pastoralists, and hunter-gatherers, with nominally significant differences in 21 tissues (Supplementary Table 2). In each tissue, hunter-gatherers had significantly lower *FADS1* levels than in agriculturalists or present-day Europeans, as would be expected from a diet containing higher levels of long-chain plasma unsaturated fatty acids (Fig. 4B). We observed a similar trend among 32 ancient Africans, indicating that this is not necessarily specific to Eurasian populations (Supplementary Fig. 7A). The variants driving these regulatory differences are in linkage disequilibrium (LD) with the functional haplotype implicated in previous evolutionary studies (Supplementary Fig. 7A; Supplementary Table 3) (Ameur *et al.*, 2012; Buckley *et al.*, 2017; Mathieson and Mathieson, 2018; Ye *et al.*, 2017). Overall, *FADS1* predicted regulation is also negatively correlated with the date of the sample (Spearman *ρ*= −0.32, *P* = 1.95 × 10^−20^, which agrees with known allele frequency trajectories (Buckley *et al.*, 2017; Mathieson and Mathieson, 2018; Ye *et al.*, 2017).

**Figure 4:**
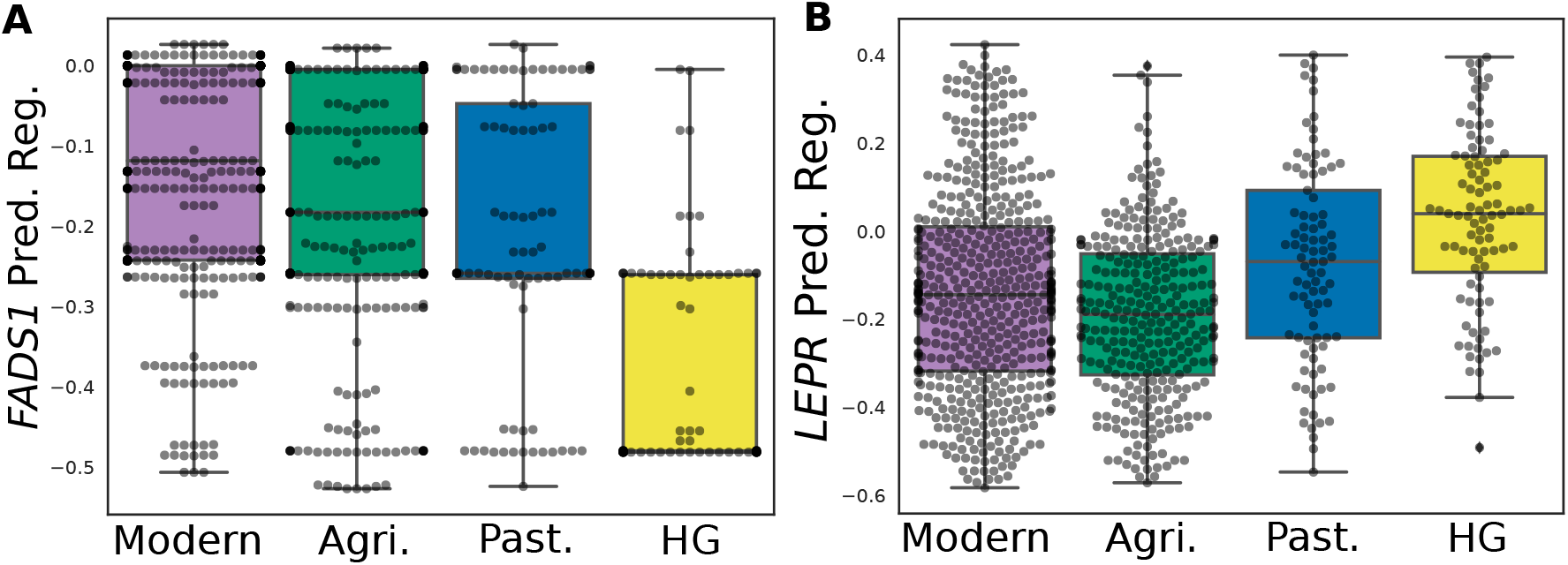
Ancient humans from different lifestyles had significant differences in regulation of key diet genes. (a) *FADS1* shows divergence in predicted regulation in Subcutaneous Adipose tissue between lifestyles (Kruskal-Wallis *P* = 5.7 × 10^−24^), as well as in eight other tissues. (b) *LEPR* regulation in Cerebellum is divergent across lifestyles (Kruskal-Wallis *P* = 3.6 × 10^−17^). Plotted with 503 present-day Europeans for comparison. Purple = Present-day Europeans, Green = agriculturalists, Blue = pastoralists, Yellow = hunter-gatherers.

Another gene in the *FADS* gene cluster, *FADS2*, functions in the same pathway as *FADS1* and is also among the 500 most diverged genes. However it shows evidence for divergent regulation in fewer tissues than *FADS1* (Supplementary Table 4), and the direction of effect is not consistent across tissues. Its presence therefore seems more likely to be due to overlap in regulatory variants with *FADS1* than to selection on *FADS2* regulation specifically. Our results further support the relevance of lifestyle differences between ancient populations in selection on the *FADS* locus and highlights the potential importance of regulatory changes of *FADS1* in dietary adaptation in Eurasians.

Among the putative diet adaptation genes, *GPX1*, an antitoxin selenoprotein, and *SLC22A5*, a transporter responsible for recycling and uptake of carnitine (Console *et al.*, 2018).(Supplementary Table 4; Supplementary Fig. 8) were also divergently regulated. The *GPX1* locus has experienced selective sweeps related to environmental selenium levels (Engelken *et al.*, 2016; White *et al.*, 2015), and has been implicated in response to viral infections (Guillin *et al.*, 2019). Carnitine plays an important role in the transport of certain long-chain fatty acids to the mitochondria for energy production; thus, modulation of its regulation could suggest a difference in metabolism related to variation in the energy demands of different lifestyles. Both selenium and carnitine levels differ in the likely primary diets of the ancient populations considered here (Flanagan *et al.*, 2010; Mann, 2018), suggesting that both as potential targets of local adaptation.

Though it was not on the list of putative diet-related adaptation genes, *LEPR* has been suggested as the driver of nearby signatures of selection due to its function in appetite and cold tolerance (Hancock *et al.*, 2008; Luca *et al.*, 2010; Voight *et al.*, 2006). *LEPR* was divergently regulated between lifestyle groups in the cerebellum (Fig. 4B) (the only brain tissue with a model for *LEPR*), both adipose tissues, and several other tissues. It was consistently predicted to be downregulated in agriculturalists compared to the other two groups in each tissue (Supplementary Table 5). Leptin is a hormone produced by adipose cells that suppresses appetite (Barrios-Correa *et al.*, 2018), so this supports a possible connection between appetite regulation and the observed signatures of selection. This is particularly relevant to modern populations given the association of decreased *LEPR* function with obesity and metabolic disorders (Dehghani *et al.*, 2018; Farooqi *et al.*, 2007).

Overall, these analyses suggest that recent regulatory changes made a substantial contribution to adaption to diet. More broadly, they demonstrate the potential for this method to explain observed signals of selection and to disentangle its effects on nearby genes.

### Skin pigmentation evolution was not driven by changes in gene regulation

We hypothesized that genes involved in complex phenotypes under selection in a population would exhibit systematic changes over time in their regulation. To test this, we focused on skin pigmentation, a trait that is known to have been under selection in humans in West Eurasia (Berg and Coop, 2014; Ju and Mathieson, 2020; Wilde *et al.*, 2014) and for which many of the genes involved are well-understood (Sturm and Duffy, 2012). We trained new PrediXcan models using genetic variants and gene expression in melanocytes from a diverse population (Zhang *et al.*, 2017). We were able to model 17 genes known to be involved in the melanogenesis pathway (Sturm and Duffy, 2012). Because skin pigmentation-associated variants changed in frequency over time, we applied these models to a time series of 2999 ancient Europeans dated between 38,052 yBP and 150 yBP, as well as 503 present-day Europeans from the 1000 Genomes Project and tested for systematic changes over time in predicted regulation.

Skin pigmentation genes are not enriched for differential regulation compared to all 6923 genes modeled in melanocytes (K-S Test *P* = 0.53; Fig. 5A). Predicted regulation showed a nominally significant linear relationship with time for only four skin pigmentation genes (Table 1), and only one (*TYR*) remained significant after genomic control.

**Figure 5:**
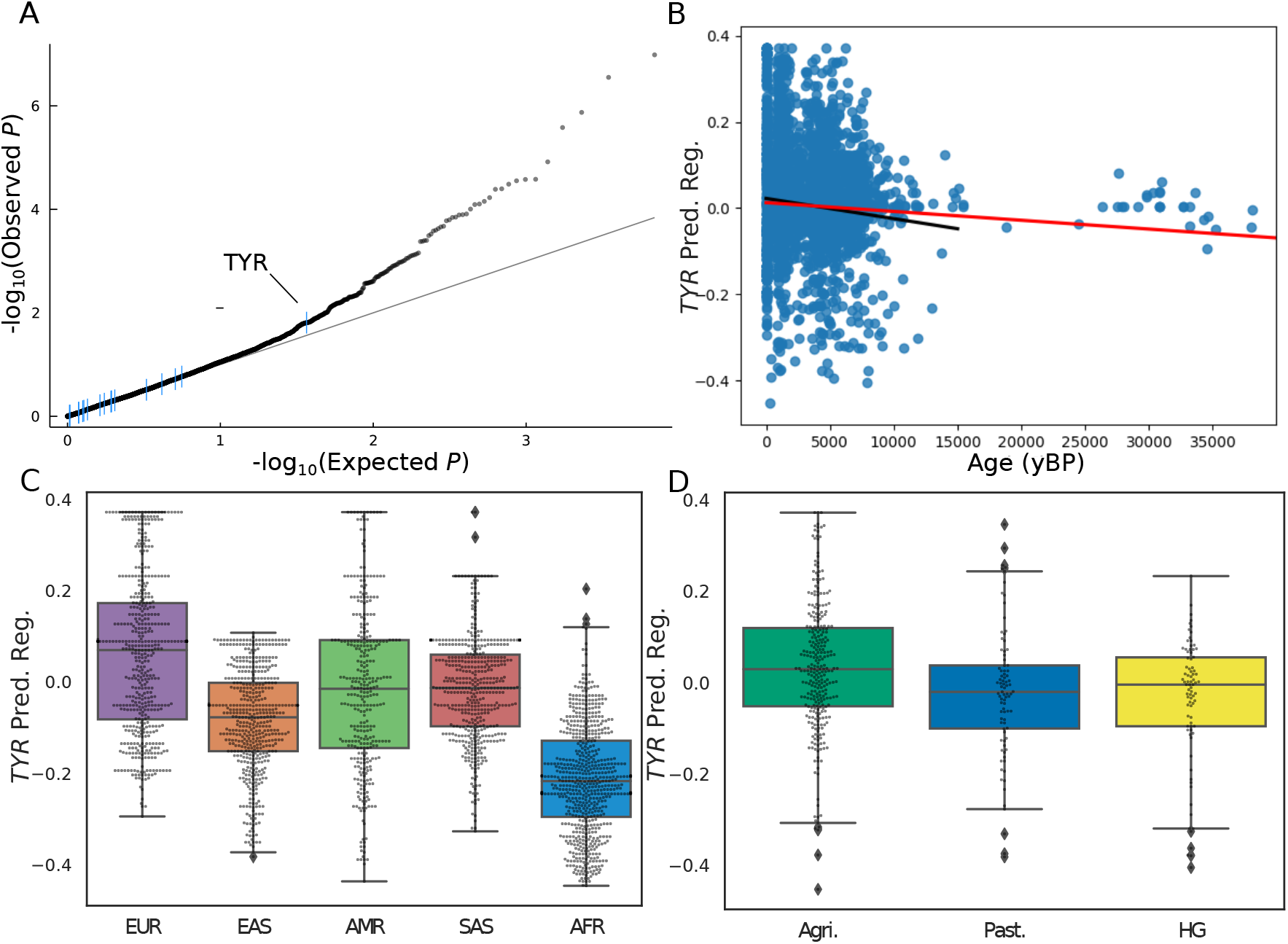
Most skin pigmentation genes show little change in regulation in the last 38,000 years in Europeans. (a) QQ plot with *P*-values from linear regressions of date vs. predicted regulation for all modeled genes in melanocytes (Methods). The 17 skin pigmentation are highlighted in blue. (b) Predicted regulation of *TYR* increases over time in Europeans. The red line shows a regression calculated over all individuals, and the black regression line was calculated only over individuals <15,000 yBP. (c) *TYR* predicted regulation in present-day 1kG populations, separated by continent of ancestry. (d) *TYR* predicted regulation in ancient Eurasians, split by lifestyle. Green = agriculturalists, Blue = pastoralists, Yellow = hunter-gatherers.

We predict that *TYR*’s expression increased over time (Fig. 5B) and is higher in non-African (particularly European) populations compared to African populations (Fig. 5C), and in (more recent) agriculturalist populations compared to hunter-gatherers (Fig. 5D). *TYR* encodes an enzyme important for one of the earliest steps of the melanogensis pathway and loss-of-function mutations cause albinism (Ghodsinejad Kalahroudi *et al.*, 2014; Norman *et al.*, 2017). It is therefore surprising that increased expression would be driven by selection for decreased pigmentation. One possibility is that increased expression due to gene regulatory variants compensates for the increase in frequency in Europeans of an activity-reducing coding variant (rs1042602) in *TYR* (Wilde *et al.*, 2014). Selection on pigmentation could favor the coding variant, while the maintenance of other functions of the gene could require increased expression. Supporting this, rs1042602 has a positive weight in the fitted PrediXcan model showing that it in fact is associated with increased expression.

Finally, we were unable to build accurate PrediXcan models for many known pigmentation genes, including those with known selected coding changes (Lamason *et al.*, 2005; Soejima and Koda, 2007), mostly because there was not enough regulatory variation nearby the genes. Overall, our results suggest that, in contrast to diet, changes in gene regulation did not play a large role in the evolution of skin pigmentation in Europe. This is consistent with observations that selection signals for pigmentation-associated variants in Europe are mostly driven by a relatively small number large-effect, coding variants despite the polygenic nature of the phenotype Ju and Mathieson (2020).

## Discussion

In this study, we adapted the PrediXcan approach for modeling the genetic component of tissue-specific gene regulation and applied it to hundreds of low-coverage ancient DNA samples from individuals from three different lifestyles and to a ~38,000-year transect of ancient Europeans. Our simulations and evaluations suggest that models of gene regulation for thousands of genes retain utility even when variant data are limited, as long as the models are trained for the specific application and their limitations properly taken into account. This is encouraging for the expansion of the PrediXcan approach to other contexts in which different variants were assayed than those used to train the original PrediXcan models. As more accurate methods are developed, it will be important to keep this aspect of their performance in mind.

Here, we found that over 5,000 genes showed evidence for divergent regulation among ancient hunter-gatherers, pastoralists, and agriculturalists in at least one tissue. The 500 genes most divergently regulated between lifestyles were enriched for metabolic and immune processes, indicating that altered gene regulation has shaped these functions during recent human evolution. Focusing on genes involved in diet, we find enrichment for divergent regulation in genes with nearby signals of recent selection, suggesting that changes in gene regulation may play a substantial role in adaptation to changes in diet.

Second, we trained new prediction models in melanocytes to analyze changes in the regulation of skin pigmentation genes in a time transect of ancient and present-day Europeans spanning 38,000 years. In contrast to genes associated with diet, we found that most genes we modeled show little to no systematic change in regulation over time, suggesting that selection on skin pigmentation mostly operated on a few large-effect coding variants. The exception, *TYR*, is predicted to have been up-regulated over time, which is contrary (with repsect to the trait) to the effects of a known coding variant in the gene and the predicted effects of gene expression on the trait itself (Chaki *et al.*, 2011; Wilde *et al.*, 2014). However, the increased expression in Europeans may be a response to the increase in frequency of a coding variant (rs1042602) that decreases activity. These results underscore the wide variety of adaptive mechanisms in recent human evolution, and the ability of ancient DNA to illuminate these mechanisms. The other skin pigmentation genes that show nominal changes in predicted regulation over time, *MITF* and *TRPM1*, are closely linked to *TYR* in the melanogenesis pathway, with *MITF* regulating both *TYR* and *TRPM1* (D’Mello *et al.*, 2016). Further analysis of the predicted perturbations of those relationships is needed to better understand the phenotypic consequences of these changes.

There are a several caveats to consider when interpreting these PrediXcan results. Previous work has demonstrated that, while there are some decreases in accuracy, the approach maintains utility when applied to non-European present-day populations and to archaic hominins (Colbran *et al.*, 2019; Petty *et al.*, 2019). Furthermore, the ancient Eurasian individuals considered here are less diverged from the GTEx cohort used for training than in these previous applications. However, due to the low coverage of the aDNA data and the focus on commonly assayed variants, there are many regulatory effects that these models do not capture. In addition, the models do not capture the effects of environment (both direct and indirect) on gene expression. Therefore, while differences in predicted regulation do not necessarily indicate a change in transcript expression levels, they do the identify change in the genetic architecture of a gene’s regulation. Our approach is therefore complementary to experimental assays of the regulatory effects of ancient genomic variants in present-day human cells (Weiss *et al.*, 2021), and such approaches could be used to test our computational predictions. Another major limitation is that we are only able to draw conclusions about genes with sufficient expression and nearby present-day common variation. Finally, we have not developed a formal test for selection on gene regulation. While we have in some cases been able to link regulatory variation to signals of selection based on genomic data, most of the differences we observe were likely the result of genetic drift. Developing tests for selection on gene regulation that consider aDNA remains an important area for future work.

Despite these limitations, we demonstrate the utility of considering regulatory effects of variants in combination in ancient individuals. In particular, we show that changes in gene regulation were essential to many, but not all, recent human adaptations. The frequent occurrence of metabolic and immune genes among the most divergently regulated genes between ancient lifestyles underscores the contribution of gene regulation to adaptation to the substantial changes in lifestyle that the shift from nomadic hunting and gathering to stationary farming had on humans. Our targeted analysis of diet genes with evidence of results adaptive evolution further suggests that adapting to diets with different nutrient and fat compositions required population-level shifts in the regulation of many metabolic genes. In contrast, the lack of consistent gene regulatory changes in skin pigmentation genes suggests that adaptation in this trait was mainly mediated by coding variants.

Lifestyle and sun exposure are not the only variables that differ among the ancient humans with genetic information, and more diverse aDNA data are rapidly becoming available. Therefore, extending this analysis to ancient individuals across other evolutionary shifts will promising. It will also be informative to expand studies into non-European populations, both ancient and present-day, to learn when gene regulatory shifts are unique to specific populations or shared.

Overall, this study demonstrates the power of focusing evolutionary analyses on combinations of variants with established relationships to molecular phenotypes. Our approach is well-positioned to use the increasing availability of present-day and ancient genome data to provide both mechanistic explanations of selection signals and to generate hypothesis about phenotypic differences between ancient and present-day groups. While this study focused on gene regulatory shifts in response to changes in lifestyle and temporal shifts in regulation of skin pigmentation genes, similar methods could be applied in many other questions and sets of ancient samples. Given the importance of gene regulation in recent evolution, this is a necessary step in identifying and interpreting candidate regions that have been shaped by recent human evolution. Further analyses using this approach will contribute to understanding the genome’s response to large-scale environmental changes and the influence of these changes on humans today.

## Methods

### Ancient genotype and lifestyle data

For the lifestyle analyses, we obtained ancient human genotypes from a set compiled and analyzed by the Allen Ancient DNA Resource (v42.4; accessed March 1, 2020), then lifted them over the Genome Build hg38 using liftOverPlink. We filtered out samples that did not pass their QC procedure and ranked remaining samples by genotype count (i.e., the number of variants with a genotype call). We also filtered samples by their continent of origin, and primarily focused on 490 ancient Eurasians. (A FADS1 analysis additionally considered 32 ancient Africans.) For a present-day comparison, we used genome for 503 European samples from the 1000 Genomes Project (The 1000 Genomes Project Consortium, 2015).

We manually assigned ancient samples to lifestyle groups by literature review based on archaeological information about the site and previous research about the associated culture. More specifically, we used lifestyles as assigned by the original publication of the sample where available. We then propagated those lifestyle labels to other samples based on the associated culture (again, as assigned by the original publication), then conducted a further literature review to match any unassigned cultures to a lifestyle based on similarity to those already matched. Samples were removed from consideration when there was not enough lifestyle-related evidence to make a call. The distinction between pastoral and agricultural groups was often difficult, and when there was ambiguity the groups were preferentially assigned to the agricultural category (Supplemental File S1).

### Adapting PrediXcan for aDNA

#### Final models for aDNA-based gene regulation prediction

The set of models used for all lifestyle analyses were trained on whole genome sequencing and RNA-seq data from GTEx v8 for 49 tissues using 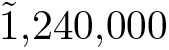 variants that were genotyped by first enriching for the targeted variants (“1240k set”) (Fu *et al.*, 2015; Haak *et al.*, 2015). For each tissue, we considered only models that explained a significant amount of variance (FDR < 0.05, *r*^2^ > 0.01). In addition, we further required that each 1240k-trained model maintain high correlations with the original GTEx model (*r* > 0.5) over all 2504 1kG individuals. All LD calculations for variants in all 1kG Populations were made using LDLink (Machiela and Chanock, 2015).

The set of models used to study skin pigmentation were trained on genotype and RNA-seq data collected from melanocytes from 106 male skin samples (Zhang *et al.*, 2017). We imputed all genotypes to 1000 Genomes using the NIH TOPMed server (Das *et al.*, 2016) with the following settings: ref: 1kG Phase 3 v5; pop = other/mixed; rsq filter 0.001; phasing = eagle v2.4. We filtered genes to those with measured expression in at least 10 samples, with RSEM > 0.5 and > 6 reads, then each gene was inverse quantile normalized to a standard normal distribution across samples. We then corrected for ancestry using the first 3 principle components and 10 PEER factors. We trained the PrediXcan models using only ~1,240,000 SNPs that were genotyped by first enriching for those targeted SNPs (“1240k set”) (Fu *et al.*, 2015; Haak *et al.*, 2015), and included any gene for which the model was able to explain a nominally significant amount of variance in the observed expression (*P* < 0.05). We focused on a set of 17 genes (Sturm and Duffy, 2012) involved in skin pigmentation for which we were able to build models.

We abbreviate the 49 GTEx tissues considered as follows: Adipose - Subcutaneous: ADPS, Adipose - Visceral Omentum: ABPV, Adrenal Gland: ADRNLG, Artery - Aorta: ARTA, Artery - Coronary: ARTC, Artery - Tibial: ARTT, Brain - Amygdala: BRNAMY, Brain - Anterior Cingulate Cortex: BRNACC, Brain - Caudate: BRNCDT, Brain - Cerebellar Hemisphere: BRNCHB, Brain - Cerebellum: BRNCHA, Brain - Cortex: BRNCTX, Brain - Frontal Cortex: BRNFCTX, Brain - Hippocampus: BRNHPP, Brain - Hypothalamus: BRNHPT, Brain - Nucleus Accumbens basal ganglia: BRNNCC, Brain - putamen basal ganglia: BRNPTM, Brain- Spinal Cord Cervical C-1: BRNSPN, Brain- Substantia Nigra: BRNSN, Breast: BREAST, Cells - Transformed Fibroblasts: FIBS, Colon - Sigmoid: CLNS, Colon - Transverse: CLNT, Esophagus - Gastroesophageal Junction: ESPGJ, Esophagus - Mucosa: ESPMC, Esophagus - Muscularis: ESPMS, Heart - Atrial Appendage: HRTAA, Heart - Left Ventricle: HRTLV, Kidney Cortex: KDNY, Liver: LIVER, Lung: LUNG, Minor Salivary Gland: MNRSG, Cells- EBV-transformed Lymphocytes: LYMPH, Ovary: OVARY, Pancreas: PNCS, Pituitary: PTTY, Prostate: PRSTT, Skeletal Muscle: MSCSK, Skin - Not sun-exposed: SKINNS, Skin - Sun-exposed: SKINS, Small Intestine: SMINT, Spleen: SPLEEN, Stomach: STMCH, Testis: TESTIS, Thyroid: THYROID, Tibial Nerve: NERVET, Uterus: UTERUS, Vagina: VAGINA, Whole Blood: WHLBLD.

#### Evaluating strategies for applying PrediXcan to aDNA

To evaluate the performance of different strategies for training PrediXcan regulation prediction models and applying them to aDNA, we carried out several simulations. In the random simulations, for each percentage missing threshold, we randomly selected 20 European individuals from 1kG (The 1000 Genomes Project Consortium, 2015), then randomly removed that percentage of genotype calls from their genomes before applying PrediXcan models to the simulated genomes (Supplementary Fig. 1). For each downsampled genome, we calculated a Spearman correlation between the predicted regulation of each gene in four tissues for the downsampled vs. the full genome. Thus, each box in Supplementary Fig. 2A has 80 (20 × 4) points. We then calculated the Spearman correlation between the median correlation between downsampled and full model predictions for each threshold and the percentage of variants missing at that threshold.

We also simulated missing data by matching patterns of missing variants from aDNA samples (Supplementary Fig. 1B). We used 3383 ancient human samples compiled and made available by the Allen Ancient DNA Resource on March 1, 2020 (v42.4). We selected three random Europeans from 1kG, then for each ancient sample we created three matching masked genomes that were missing exactly the same variants. For each masked genome, we calculated the Spearman correlation between the predicted regulation of each gene in all four tissues for the masked vs. the full genome (i.e. one correlation per individual).

We also evaluated three different sets of variants for training PrediXcan models. The “full set” consisted of all variable sites identified in GTEx v8 (this included both single nucleotide variants and short indels in hg38 coordinates). The “1240k set” was formed by intersecting the full set with the variants genotyped on the 1240k chip, which totalled 714,959 variants after lifting them over to hg38. Lastly, we assembled the “top600k set” of variants, which is a subset of the 1240k set with high “support”. We calculated the “support” for each variant over *N* aDNA samples as 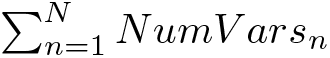, where *NumV ars* is the number of variants successfully called in sample *n*. In other words, support for a variant is the number of samples in which that variant was successfully genotyped, weighted by the quality (i.e., number of genotyped variants) of the sample. A variant can therefore obtain a high support either by being genotyped in many low-quality samples, or in fewer high-quality samples. We ranked the variants by their support. We identified the top 600k variants, and for the purposes of simulating the behaviour of models when applied to incomplete data, we also considered the top 500k variants with the highest support (“top500k”; N = 499,666). For each set of variants, we trained a set of models and created a set of 1kG genomes masked to only include those variants (Fig. 1A). We assessed the performance of combinations of models and genomes by calculating the correlation of predictions made by each model-genome pair with predictions made by the Full models on the Full 1kG Genomes (i.e. one correlation was calculated per individual 1kG sample for each pair).

### Identifying divergent gene regulation between ancient lifestyles

To identify genes with evidence for divergence in predicted gene regulation between the three lifestyle groups, we applied a Kruskal-Wallis test for the predictions of each gene model over individuals from each group. We accounted for multiple testing with a Bonferroni correction within each tissue. Genes passing the correction in at least one tissue are said to show evidence for a significant difference in regulation. To further isolate the genes that are the most likely to be diverged due to selection rather than drift, used genomic control to correct for population stratification by calculating an inflation factor *λ* and recalculating p-values based on the distribution of *χ*^2^/*λ* (Devlin and Roeder, 1999). To focus our discussion on the genes with the strongest evidence for divergence, we sorted all models by GC-corrected *P*-value and identified the top 500 unique genes (corresponding to 1236 models), which corresponded to those with at least one model with a GC-corrected *P* < 3.46 × 10^−3^ and FDR=0.586.

### Gene set enrichment among diverged genes

To conduct functional enrichment analyses on the top 500 most diverged genes, we tested for GO annotation over-representation using WebGestalt with default parameters (Liao *et al.*, 2019) Specifically, we compared the biological process GO terms among the 500 most diverged genes versus all genes with a model in at least one tissue. We also tested for enrichment of several other gene sets of interest: 1) genes whose expression in particular tissues is under stabilizing selection across 17 mammalian species (Chen *et al.*, 2018); genes that are intolerant to loss-of-function variants in their protein products (called if the upper bound of the 95% confidence interval of the observed/expected ratio is lower than 0.35) (Lek *et al.*, 2016); 3) housekeeping genes that show consistent expression across tissues (Eisenberg and Levanon, 2013); and 4) a set genes encoding virus interacting proteins (Enard *et al.*, 2016). We calculated an odds ratio for each, and used a Fisher’s exact test to determine significance. For the genes under stabilizing selection on gene expression, we considered only those tested in that study before calculating statistics.

### Skin pigmentation time series data and analysis

We obtained ancient human genome data from the Allen Ancient DNA Resource (v44.3; accessed February 8, 2021). We filtered for individual human samples from Europe (west of 59° East), and in the case of duplicate individuals chose the sample with the highest average coverage. We filled in missing dosages using the mean dosage across the other samples. This resulted in 2999 ancient Europeans, to which we added 503 European samples from the 1000 Genomes Project (The 1000 Genomes Project Consortium, 2015) to construct a time series ranging from 38,052 yBP to present (31 samples were older than 15,000 yBP).

To identify genes which showed a systematic change in regulation over time, we obtained predicted regulation values for each gene in each individual using the melanocyte PrediXcan models. We then regressed the predicted regulation on the date of the sample using a linear regression framework, including the first 10 principle components to correct for ancestry. We further controlled for population stratification using genomic control (Devlin and Roeder, 1999), and identified the skin pigmentation genes for which the effect size of date was significant (corrected *P* < 0.05). We additionally compared the predicted regulation of *TYR* in all 2504 individuals from the 1000 Genomes Project (The 1000 Genomes Project Consortium, 2015), separated by continental ancestry.

## Data and Code Availability

All data and scripts are available on Github at https://github.com/colbrall/ancient_human_predixcan and https://github.com/colbrall/skin_pigmentation_regulation

## Acknowledgements

We would like to thank Eric Gamazon for discussions and help with data acquisition, as well as Dan Zhou for help in debugging the PrediXcan training pipeline. We would also like to thank current and former members of the Capra and Mathieson labs for useful discussions of the project and for providing comments on the figures and manuscript.

L.L.C. was funded by NIH grant T32GM080178 to Vanderbilt University and T32HG009495 to the University of Pennsylvania. I.M was funded by NIH award R35GM133708. J.A.C. was funded by NIH award R01GM115836 and R35GM127087, and the Burroughs Wellcome Fund.

This work was conducted in part using the resources of the Advanced Computing Center for Research and Education at Vanderbilt University, Nashville, TN. The research is solely the responsibility of the authors and does not necessarily represent the official views of Vanderbilt University Medical Center, the National Institutes of Health, or other funding agencies.

## Author Contributions

L.L.C., I.M. and J.A.C. designed the experiments and wrote the manuscript. M.R.J. designed the simulation experiments and conducted pilot simulation analyses. L.L.C conducted all other experiments. All authors edited and approved the final manuscript.

## Supplementary Materials

### Evaluating the robustness of gene regulation prediction models to missing data

There are several challenges involved in adapting PrediXcan to be applied to ancient DNA. First, ancient individuals may be genetically diverged from the populations used to train the prediction models. Recent evaluations of the portability of these models across modern human populations have shown that while their accuracy decreases when applied across human populations, most models maintain substantial predictive ability that enables the discovery of meaningful biological associationsOkoro *et al.* (2021); Petty *et al.* (2019). We have also demonstrated that this approach can be applied to archaic hominin genomesColbran *et al.* (2019). The ancient Eurasian individuals we analyze here are less genetically diverged from the training population than modern African-ancestry individuals or archaic hominins. In addition, we stress that these models do not aim to predict gene expression itself, but rather to quantify the common variant mediated component of gene regulation, and thus, they provide an means to detect shifts in regulatory architecture.

Second, available aDNA data varies in coverage, depth, and quality. This creates a trade-off between number of individuals available for analysis and the quality of their genotyping. In addition, many other potential applications of PrediXcan involve populations that differ from the training populations. In these situations, even if the populations are of similar ancestry, all the variants the models use to predict gene expression may not be assayed in the population of interest. Therefore it is of great interest to understand how PrediXcan behaves under these conditions with varying levels of missing variant data, and develop ways to optimize its performance in such cases.

To better understand and quantify these patterns, we first trained PrediXcan models on expression data and all available variants in GTEx v8 (“Full models”). We then applied these prediction models to all variants and downsampled sets of variants from individuals with whole-genome sequencing from the 1000 Genomes Project (1kG)The 1000 Genomes Project Consortium (2015). This enabled us to evaluate how predictions change with missing variant data. We selected nine thresholds for percentage of missing SNPs (5%-45% missing) and downsampled 20 random European individuals per threshold (Supplementary Fig. 1A). The agreement between the predictions on downsampled genomes and full genomes was strongly correlated with the percentage of SNPs missing (Supplementary Fig. 2B). However, Spearman correlations were above 0.75 for all comparisons, even when genomes were missing as many as 45% of their SNPs. This suggests that model accuracy can be maintained even at relatively high rates of missingness, likely due to LD between variants.

While this is encouraging, missing SNPs may not be randomly distributed throughout the genome in some applications. To evaluate how biases in missingness across the genome could affect these results, we repeated the comparison described above. However, instead of randomly downsampling SNPs, we matched the patterns of missingness to the dataset (Fig. 1B) of particular interest to us: 3383 aDNA samples, with widely varying numbers of missing SNPs (Supplementary Fig. 2C). Overall, the correlations were much lower (median Spearman *ρ*=0.39; Fig. 1B). This is unsurprising given that the aDNA samples were obtained using targeted capture, while the model training data was based on whole genome sequencing. At most, the aDNA samples had 714,959 SNPs, while the training data had over 5 million (i.e. 87% missing). This indicates that, while models can tolerate a fair amount of missing data, the missingness caused by a mismatch between genotyped SNPs and training SNPs is likely to substantially decrease prediction accuracy.

We next evaluated whether imputation performance could be improved in our aDNA application by customizing the training data to contain only variants that will be available in the application data. This step ensures that any variants used that are not assayed in the application data are not used in model training. Thus, we retrained PrediXcan models using GTEx v8, but only considered the SNPs present in the 1240k capture set that is commonly used in aDNA studies (“1240k set”; Supplementary Fig. 1B). This resulted in 714,959 variants for model training (the number that were successfully lifted over to the hg38 genome build), as opposed to the 5,310,489 variants available for the full models based on whole genomes. Because most of the genotyped aDNA samples tend to be low-coverage (Supplementary Fig. 2C), we also tested the use-case where we chose the SNPs in the dataset most likely to provide information. To do this, we ranked SNPs by the number of samples in the 3383 ancient samples used above with genotype calls, and weighted that count by the overall coverage of those samples. We then chose the 600,000 SNPs with the best ranking (“top600k set”), thereby prioritizing SNPs that were frequently present in the best-quality samples.

Training models on fewer SNPs resulted in a small decrease in the variance explained in gene expression for some genes (Fig. 3), and fewer significant models were constructed in each tissue (7184, 6587, and 5196 significant models for Full, 1240k, and top600k respectively in Whole Blood). This also resulted in inconsistent predictions when models were applied to the same individuals (Fig. 4). This behaviour is caused by the decrease in available SNPs for modelling, which is reflected in the smaller models built in the 1240k and top600k sets (Fig. 5A,C). While many genes continued to be successfully modelled by the targeted models, restricting to fewer SNPs resulted in a loss of genes for which the Full models had a lower *r*^2^.

While these results demonstrate that using more SNPs during training results in more numerous and accurate models, this does not take into account the presence of missing data in the genomes that will be used for predictions. Therefore the outstanding question is whether model performance is more consistent when trained on fewer SNPs without a large missing data percentage, or if it is better to include more SNPs during training, but allow more missing data during application. To answer this, we applied our targeted models described above to 1kG genomes downsampled to match the various sets of SNPs used during model training and compared their agreement with the full models applied to the full genomes. For the top600k models, we further downsampled the application set to 500k SNPs using the same methodology.

In line with our initial findings, we found that all models lost consistency when applied to genomes with missing SNPs (Fig. 1C). The 1240k models maintained the highest agreement with the Full models when applied to incomplete data (median *ρ*=0.78, vs 0.58 and 0.41 for Full and top600k models, respectively), though they also had a larger variance in agreement. The Full models likely did worse because there was a much larger drop in the number of SNPs available compared to training, while the top600k models’ reliance on fewer SNPs may have increased their susceptibility to missing SNPs. This suggests that a balance between targeting model training for the dataset in question and allowing some missingness is the best course of action. For analyses presented in this paper, we therefore applied models targeted to the 1240k variant set to ancient individuals with relatively high coverage (above the 3rd quartile among all samples available).

## Supplementary Figures

**Figure 1:**
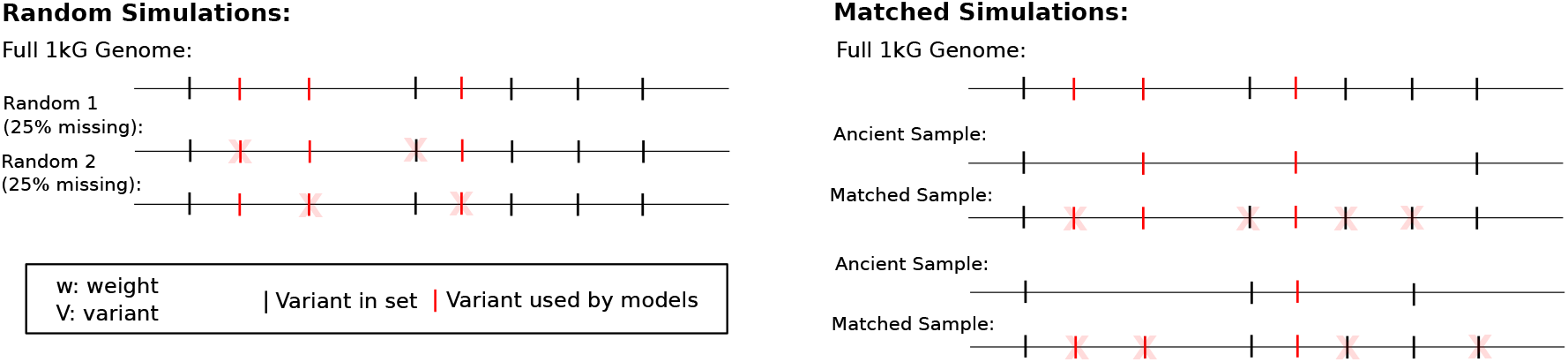
Schematic of process for generating random and matched simulated genomes. Starting from a complete genome from 1kG, for the random simulations we mask a random number of variants corresponding to the specified missing percentage. For the matched simulations, we pair a complete modern genome and an ancient genome, and mask any variants in the modern genome that are not present in the ancient one. (b) Schematic of creating targeted variants sets to train PrediXcan models.

**Figure 2:**
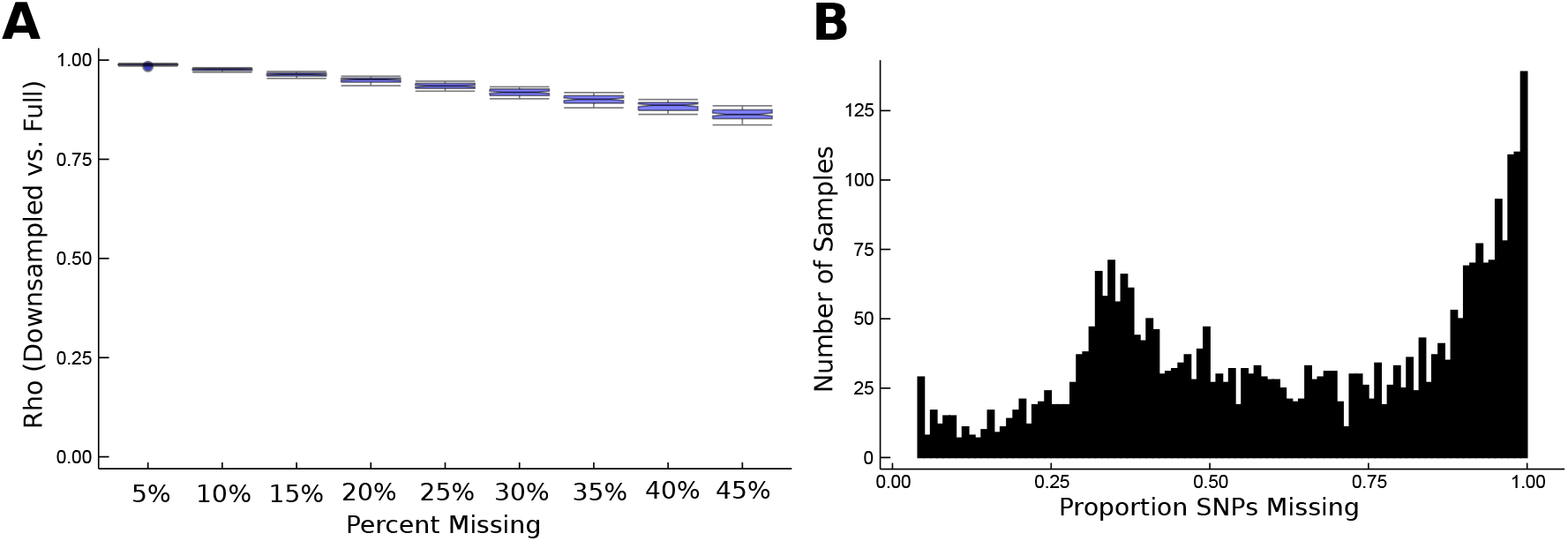
Missing variants affect PrediXcan performance. (a) The Spearman *ρ* between predictions in four tissues calculated for the complete genome vs. random simulations decreases as the percentage of missing variants increases, but remains high even with 45% of variants masked. (c) Distribution of the proportion of missing variants in the aDNA data compared to all variants included in 1kG.

**Figure 3:**
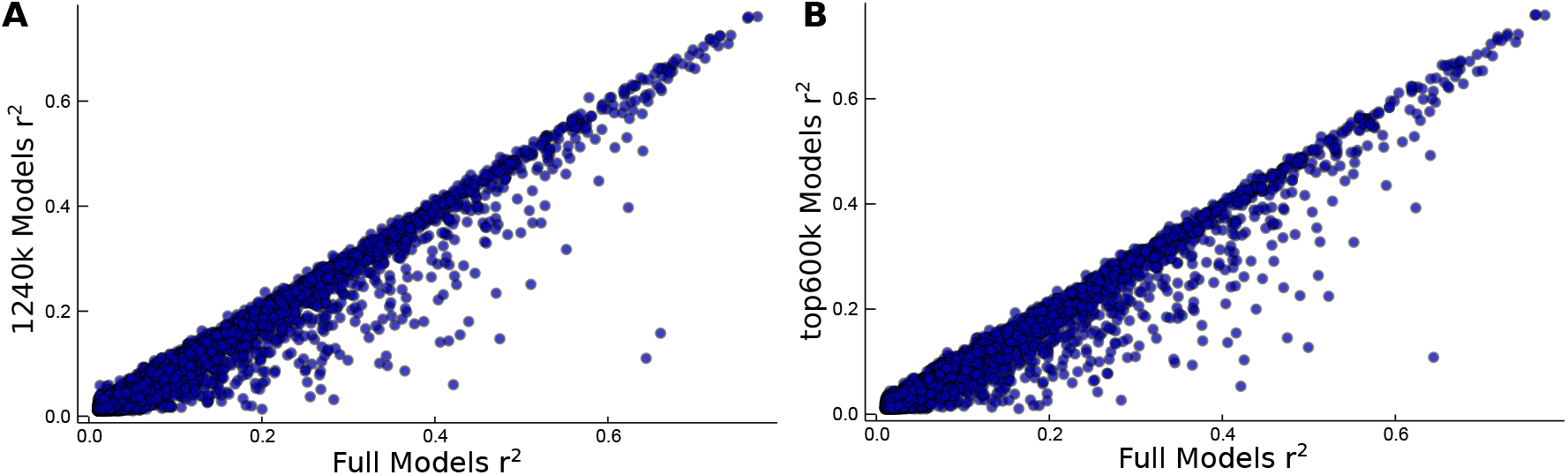
Models trained on fewer SNPs explain less variance in gene expression. Scatterplots of training *r*^2^ for (a) 1240k models and (b) top600k models vs. Full models in Whole Blood. Other tissues tested matched trends. *r*^2^ is calculated over observed vs. predicted expression.

**Figure 4:**
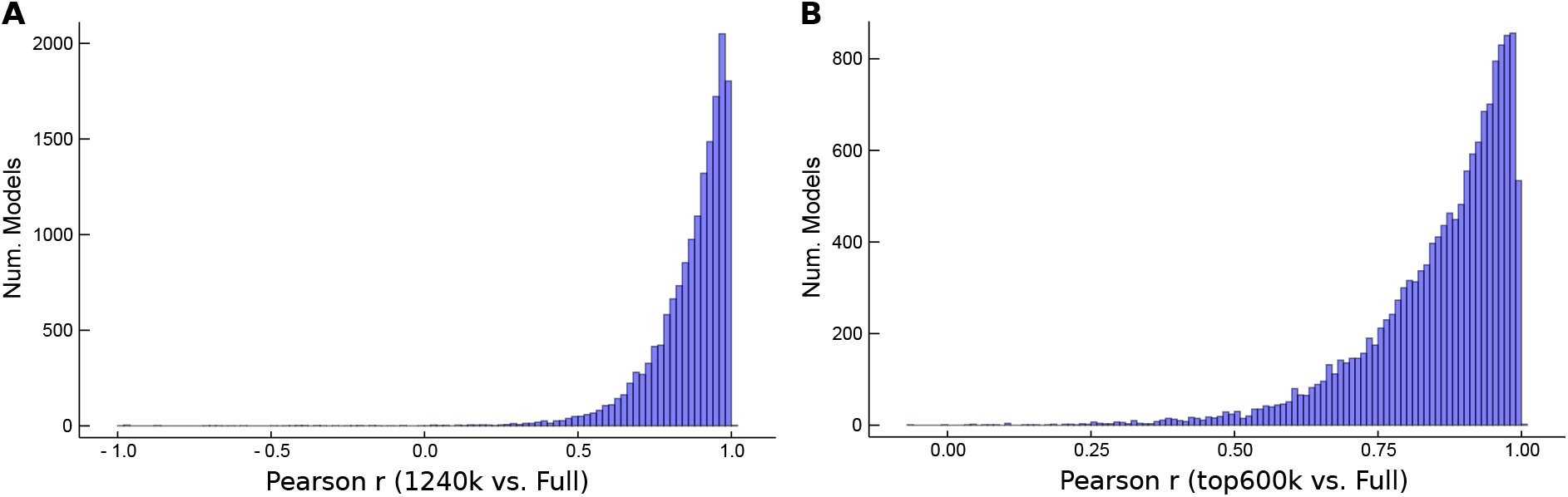
Targeted models predict consistent gene regulatory patterns. (a) Pearson r by model between Full model 1kG predictions and (a) 1240k and (b) top600k models for all genes in 4 tissues-Liver, Ovary, Whole Blood, Skeletal Muscle. Pearson correlation calculated for each model between predictions made on all 1kG individuals using Full models vs. 1240k or top600k models.

**Table 1:**
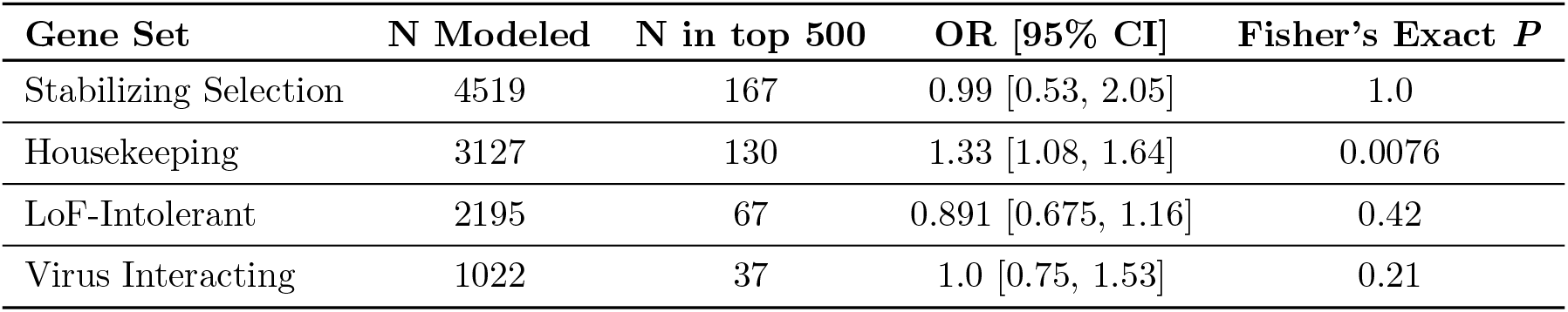
Enrichment for four gene sets of interest in the 500 most diverged genes. The odds ratio was calculated as the odds of a gene’s presence in the category given it is in the top 500 divergently regulated genes. The analysis only includes genes that were both considered for inclusion in the set and had at least one PrediXcan model. For example, genes not tested for stabilizing selection on gene expression were not considered.

| Gene Set | N Modeled | N in top 500 | OR [95% CI] | Fisher's Exact $P$ |
| --- | --- | --- | --- | --- |
| Stabilizing Selection | 4519 | 167 | 0.99 [0.53, 2.05] | 1.0 |
| Housekeeping | 3127 | 130 | 1.33 [1.08, 1.64] | 0.0076 |
| LoF-Intolerant | 2195 | 67 | 0.891 [0.675, 1.16] | 0.42 |
| Virus Interacting | 1022 | 37 | 1.0 [0.75, 1.53] | 0.21 |

**Figure 5:**
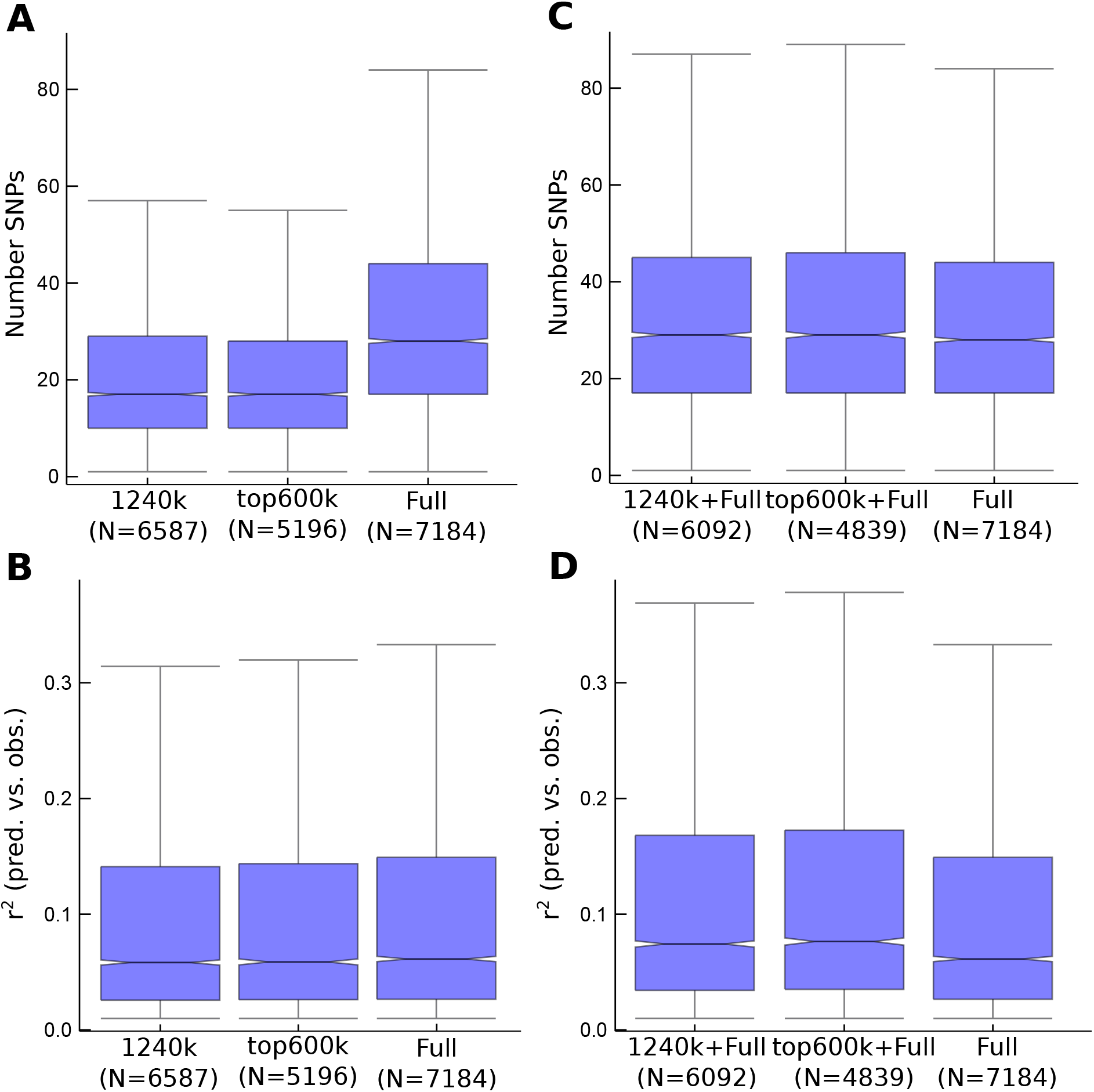
Targeted models use fewer SNPs and and succeed in modelling fewer genes. (a) Number of SNPs in each set of models for Whole Blood. (b) *r*^2^ between predicted and observed expression for each model in Whole Blood. By definition, significant models had to have *r*^2^ > 0.01 and have a within-tissue FDR < 0.05. (c) Number of SNPs and (d) *r*^2^ in models shared (identified as significant) in both the Full set and either the 1240k or top600k Set (metrics plotted are those from the Full set). Full set replotted on all plots for comparison. Other tissues tested showed similar trends.

**Figure 6:**
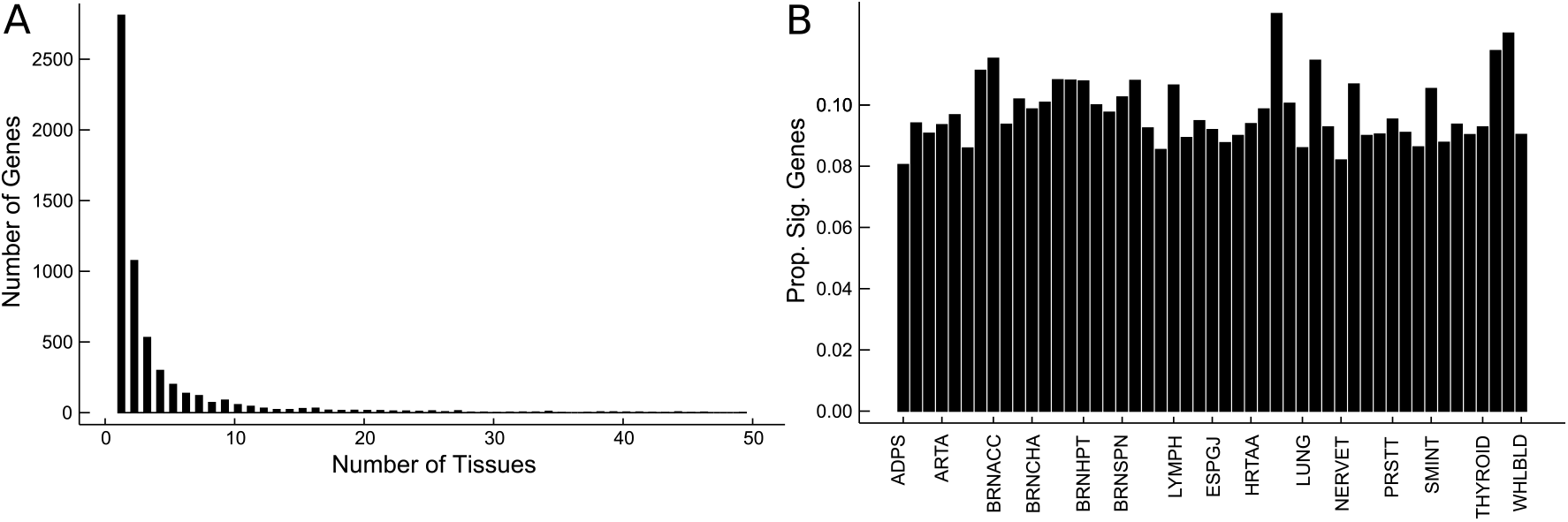
Thousands of genes are divergently regulated between lifestyle groups. (A) Distribution of the number of tissues in which each gene (N = 5760) is significantly different between groups. (B) The proportion of significant models out of all models in a tissue. Tissues are in alphabetical order: **Adipose_Subcutaneous**, Adipose_Visceral_Omentum, Adrenal_Gland, **Artery_Aorta**, Artery_Coronary, Artery_Tibial, Brain_Amygdala, **Brain_Anterior_cingulate_cortex**, Brain_Caudate_basal_ganglia, Brain_Cerebellar_Hemisphere, **Brain_Cerebellum**, Brain_Cortex, Brain_Frontal_Cortex, Brain_Hippocampus, **Brain_Hypothalamus**, Brain_Nucleus_accumbens_basal_ganglia, Brain_Putamen_basal_ganglia, **Brain_Spinal_cord_cervical_c-1**, Brain_Substantia_nigra, Breast_Mammary_Tissue, Cells_Cultured_fibroblasts, **Cells_EBV-transformed_lymphocytes**, Colon_Sigmoid, Colon_Transverse, **Esophagus_Gastroesophageal_Junction**, Esophagus_Mucosa, Esophagus_Muscularis, **Heart_Atrial_Appendage**, Heart_Left_Ventricle, Kidney_Cortex, Liver, **Lung**, Minor_Salivary_Gland, Muscle_Skeletal, **Nerve_Tibial**, Ovary, Pancreas, Pituitary, **Prostate**, Skin_Not_Sun_Exposed_Suprapubic, Skin_Sun_Exposed_Lower_leg, **Small_Intestine_Terminal_Ileum**, Spleen, Stomach, Testis, **Thyroid**, Uterus, Vagina, **Whole_Blood**.

**Figure 7:**
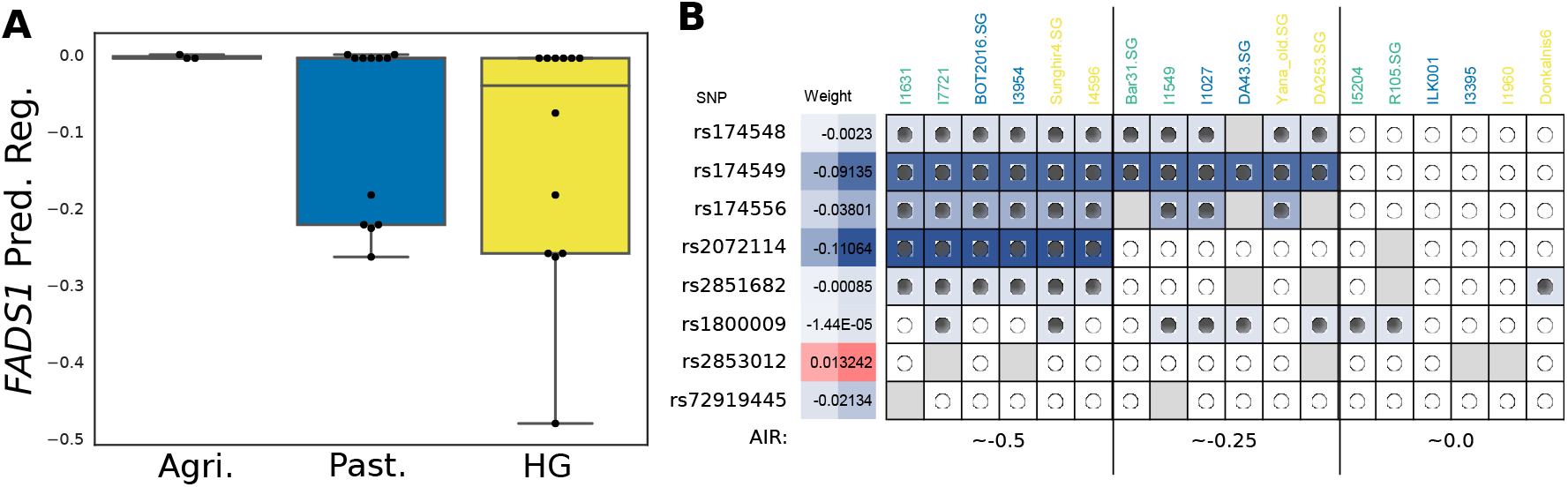
*FADS1* model variants tag a selected haplotype. (a) 27 ancient Africans follow a similar trend in expression differences to that seen in ancient Eurasian populations. (b) Breakdown of the 8 SNPs in the model of FADS1 in Adipose Subcutaneous tissue and their presence in representative ancient Eurasians across a range of predicted normalized expression values. Cells are coloured by the weight that SNP contributed to the prediction, while the circles indicate the alleles present in that individual (filled = homozygous effect, empty = homozygous reference). A grey square indicates the SNP was ungenotyped in that individual. The vast majority of ancient samples appear homozygous due to being extremely low-coverage, such that many sites are represented by only a single read.

**Figure 8:**
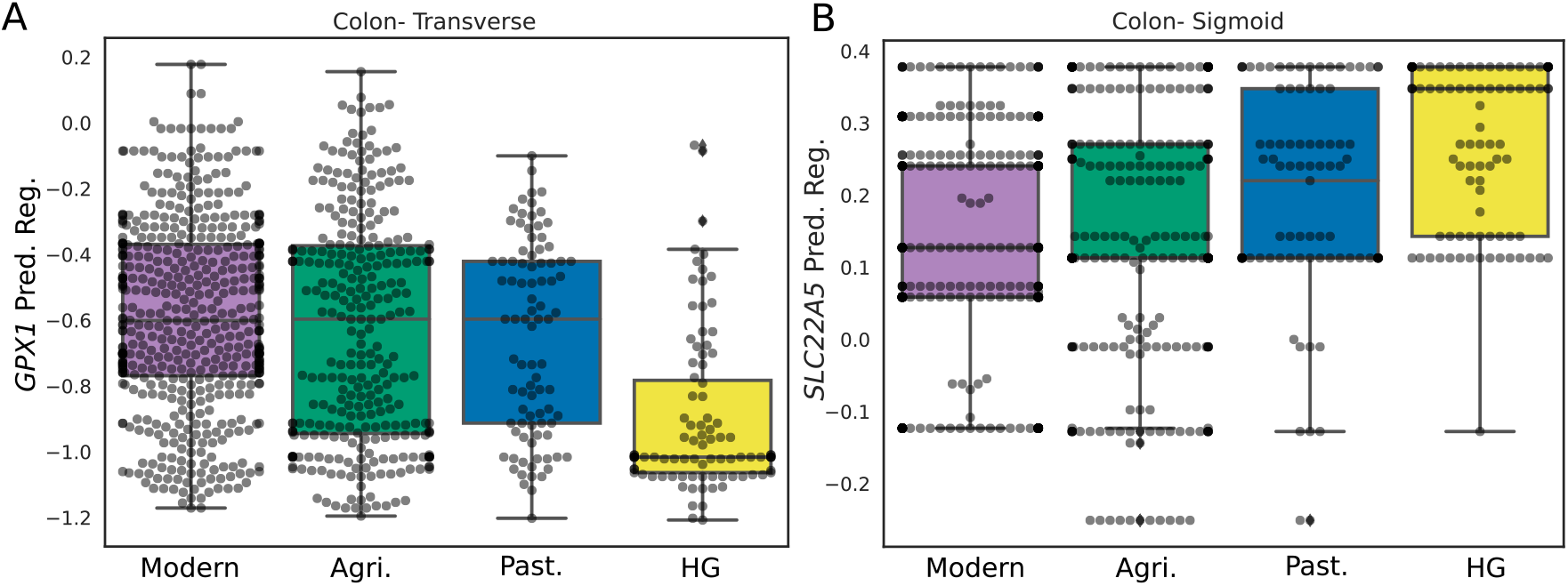
Divergently regulated genes include. (A) *GPX1* in Transverse Colon and (B) *SLC22A5* in Sigmoid Colon. Plotted with 503 modern Europeans for comparison in purple. Green = Agriculturalists, Blue = Pastoralists, Yellow = Hunter-Gatherers.

**Table 2:** *FADS1* models with divergent regulation by lifestyle. Median predicted regulation for each group*P*-value is calculated from Kruskal-Wallis test, after GC-correction. *FADS1* was modeled in an additional 9 tissues. **model *P*-value is below that corresponding to the top 500 unique genes.

| Tissue | Agri. | Past. | HG | Corrected *P* |
| --- | --- | --- | --- | --- |
| Adipose.Subcutaneous** | -0.183 | -0.259 | -0.480 | $8.28 \times 10^{-6}$ |
| Adipose.Visceral.Omentum | -0.132 | -0.158 | -0.158 | 0.0554 |
| Brain.Cerebellar.Hemisphere** | -0.882 | -1.228 | -1.422 | $1.11 \times 10^{-4}$ |
| Brain.Cerebellum** | -0.887 | -1.184 | -1.716 | $5.76 \times 10^{-5}$ |
| Brain.Frontal.Cortex** | -0.391 | -0.602 | -0.690 | $2.59 \times 10^{-9}$ |
| Brain.Putamen.basal.ganglia | -0.360 | -0.469 | -0.530 | 0.0596 |
| Cells.Cultured.fibroblasts | -0.712 | -0.768 | -1.024 | 0.0168 |
| Colon.Sigmoid** | -0.360 | -0.456 | -0.859 | $2.69 \times 10^{-5}$ |
| Esophagus.Gastroesophageal.Junction** | -0.378 | -0.426 | -0.579 | $3.06 \times 10^{-5}$ |
| Esophagus.Mucosa | -0.620 | -0.769 | -0.836 | $9.2 \times 10^{-3}$ |
| Esophagus.Muscularis | -0.319 | -0.409 | -0.506 | 0.117 |
| Heart.Atrial.Appendage | -0.225 | -0.281 | -0.382 | 0.016 |
| Heart.Left.Ventricle** | -0.169 | -0.330 | -0.924 | $2.53 \times 10^{-6}$ |
| Lung** | -0.106 | -0.154 | -0.237 | $4.2 \times 10^{-6}$ |
| Muscle.Skeletal** | -0.410 | -0.674 | -0.787 | $9.77 \times 10^{-7}$ |
| Nerve.Tibial | -0.560 | -0.748 | -0.868 | 0.0135 |
| Pancreas** | -0.452 | -0.882 | -1.414 | $1.6 \times 10^{-4}$ |
| Stomach** | -0.401 | -0.641 | -0.872 | $1.97 \times 10^{-6}$ |
| Testis** | -0.610 | -0.774 | -0.862 | $1.46 \times 10^{-3}$ |
| Thyroid** | -0.295 | -0.483 | -0.649 | $2.7 \times 10^{-5}$ |
| Whole.Blood | 0.0914 | 0.108 | 0.132 | 0.0759 |

**Table 3:**
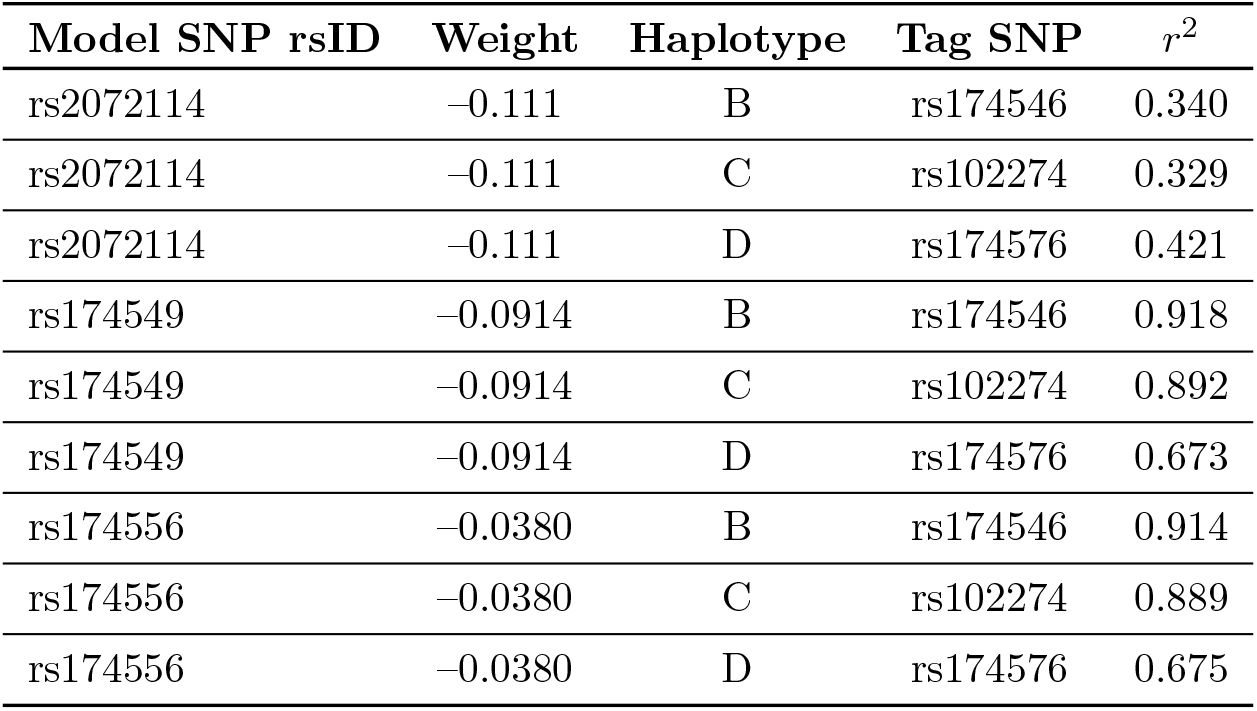
*FADS1* model SNPs LD with established haplotype. The 3 highest-weight SNPs from Subcutaneous Adipose. Haplotype labels correspond to those in Mathieson & Mathieson (2018)Mathieson and Mathieson (2018).

**Table 4:** Dietary gene models with divergent regulation by lifestyle. Median predicted regulation for each group. *P*-value is calculated from Kruskal-Wallis test, after GC-correction. *FADS1* models in Table 2 also belong in this table, but are not reprinted to avoid redundancy. **model *P*-value is below that corresponding to the top 500 unique genes.

| Gene | Tissue | Agri. | Past. | HG | P |
| --- | --- | --- | --- | --- | --- |
| HFE | Brain_Nucleus_accumbens_basal_ganglia | -0.538 | -0.426 | -0.280 | 0.0523 |
| TRIP4 | Brain_Cerebellar_Hemisphere | -0.030 | -0.067 | -0.203 | 0.0458 |
|  | Brain_Frontal_Cortex | -0.046 | -0.053 | -0.098 | 0.0868 |
|  | Skin_Not_Sun_Exposed_Suprapubic | -0.537 | -0.621 | -0.684 | 0.054 |
| FADS2 | Breast_Mammary_Tissue | -0.046 | 0.020 | 0.087 | 0.0332 |
|  | Esophagus_Muscularis | 0.255 | 0.416 | 0.454 | 0.0714 |
|  | Pancreas | 0.155 | 0.204 | 0.011 | 0.0187 |
|  | Skin_Not_Sun_Exposed_Suprapubic | 0.239 | 0.266 | 0.266 | 0.0433 |
|  | Testis** | 0.414 | 0.457 | 0.459 | 2.49e-3 |
| SLC22A4 | Cells_Cultured_fibroblasts | 0.304 | 0.360 | 0.304 | 0.0656 |
|  | Thyroid | -0.037 | -0.036 | -0.141 | 0.0329 |
| SLC22A5 | Adipose_Subcutaneous** | 0.253 | 0.253 | 0.315 | 1.94e-3 |
|  | Adipose_Visceral_Omentum | 0.054 | 0.054 | 0.054 | 0.0232 |
|  | Artery_Tibial | 0.602 | 0.736 | 0.746 | 0.0337 |
|  | Cells_Cultured_fibroblasts** | 0.644 | 0.705 | 0.832 | 0.0019 |
|  | Cells_EBV-transformed_lymphocytes | 0.182 | 0.326 | 0.366 | 0.00352 |
|  | Colon_Sigmoid** | 0.113 | 0.230 | 0.348 | 0.00324 |
|  | Esophagus_Gastroesophageal_Junction | -0.139 | -0.039 | 0.174 | 0.00498 |
|  | Esophagus_Muscularis | 0.180 | 0.255 | 0.421 | 0.00413 |
|  | Minor_Salivary_Gland | 0.228 | 0.253 | 0.262 | 0.0935 |
|  | Nerve_Tibial | 0.655 | 0.688 | 0.781 | 0.0307 |
|  | Pancreas | -0.099 | -0.074 | 0.024 | 0.0112 |
|  | Thyroid | 0.320 | 0.385 | 0.398 | 0.0368 |
|  | Whole_Blood | 0.781 | 0.910 | 1.043 | 0.0115 |
| DI2 | Nerve_Tibial | -0.239 | -0.312 | -0.389 | 0.0644 |
| AS3MT | Adipose_Visceral_Omentum | -0.143 | -0.097 | -0.495 | 0.0738 |
|  | Brain_Hypothalamus | -0.328 | -0.206 | -0.688 | 0.0923 |
|  | Heart_Atrial_Appendage | -0.243 | -0.046 | -0.449 | 0.0769 |
| GPX1 | Adrenal_Gland | -0.469 | -0.529 | -0.835 | 0.0105 |
|  | Cells_Cultured_fibroblasts | -0.691 | -0.625 | -1.115 | 0.0199 |
|  | Colon_Transverse** | -0.597 | -0.597 | -1.016 | 0.00271 |
|  | Esophagus_Mucosa | -0.372 | -0.374 | -0.663 | 4.8e-3 |
|  | Prostate | -0.317 | -0.317 | -0.572 | 0.00541 |
|  | Skin_Not_Sun_Exposed_Suprapubic | -0.379 | -0.432 | -0.712 | 0.00429 |
|  | Skin_Sun_Exposed_Lower_leg | -0.475 | -0.444 | -0.741 | 0.00913 |
|  | Small_Intestine_Terminal_Ileum | 0.054 | 0.054 | -0.100 | 0.0573 |
|  | Thyroid | -0.705 | -0.774 | -0.836 | 0.0335 |

**Table 5:** *LEPR* models with divergent regulation by lifestyle. Median predicted regulation for each group*P*-value is calculated from Kruskal-Wallis test, after GC-correction. *LEPR* was modeled in an additional 11 tissues. **model *P*-value is below that corresponding to the top 500 unique genes.

| Tissue | Agri. | Past. | HG | Corrected $P$ |
| --- | --- | --- | --- | --- |
| Adipose_Subcutaneous | -0.504 | -0.374 | -0.325 | 0.0151 |
| Adipose_Visceral_Omentum** | -0.251 | -0.0661 | -0.0555 | $6.61 \times 10^{-4}$ |
| Brain_Cerebellar_Hemisphere** | 0.00454 | 0.199 | 0.317 | $4.68 \times 10^{-4}$ |
| Brain_Cerebellum** | -0.191 | 0.0699 | 0.0395 | $2.54 \times 10^{-4}$ |
| Esophagus_Gastroesophageal_Junction** | -0.0109 | 0.0443 | 0.174 | $1.92 \times 10^{-3}$ |
| Heart_Atrial_Appendage** | -0.136 | -0.0663 | -0.0537 | $4.48 \times 10^{-4}$ |
| Testis** | -0.246 | -0.0970 | -0.0659 | $4.8 \times 10^{-5}$ |
| Whole_Blood | 0.0559 | 0.211 | 0.178 | 0.0725 |

**Table 6:** Top 20 enriched GO terms among the 500 genes most diverged between lifestyles.

| GO term | Num. Genes | Enrichment | $P$ |
| --- | --- | --- | --- |
| platelet morphogenesis | 3 | 5.04 | 0.020 |
| response to mitochondrial depolarisation | 3 | 5.04 | 0.020 |
| protein hydroxylation | 4 | 4.40 | 0.012 |
| peptidyl-proline modification | 6 | 4.18 | 0.0028 |
| oligosaccharide metabolic process | 5 | 3.86 | 0.0089 |
| response to interleukin-4 | 4 | 3.69 | 0.022 |
| ruffle organization | 5 | 3.25 | 0.018 |
| response to antineoplastic agent | 8 | 2.97 | 0.0053 |
| DNA-templated transcription, elongation | 8 | 2.72 | 0.0090 |
| cytokine metabolic process | 7 | 2.60 | 0.018 |
| type I interferon production | 8 | 2.51 | 0.014 |
| regulation of mitochondrion organization | 12 | 2.32 | 0.0057 |
| antigen processing and presentation | 11 | 2.30 | 0.0086 |
| regulation of inflammatory response | 22 | 2.10 | $8.6 \times 10^{-4}$ |
| carbohydrate derivative catabolic process | 11 | 2.07 | 0.018 |
| cellular modified amino acid metabolic process | 12 | 2.07 | 0.014 |
| positive regulation of response to external stimulus | 16 | 1.93 | 0.0092 |
| glycoprotein metabolic process | 20 | 1.83 | 0.0070 |
| positive regulation of cytokine production | 20 | 1.72 | 0.012 |
| negative regulation of intracellular signal transduction | 24 | 1.63 | 0.013 |

**Table 7:**
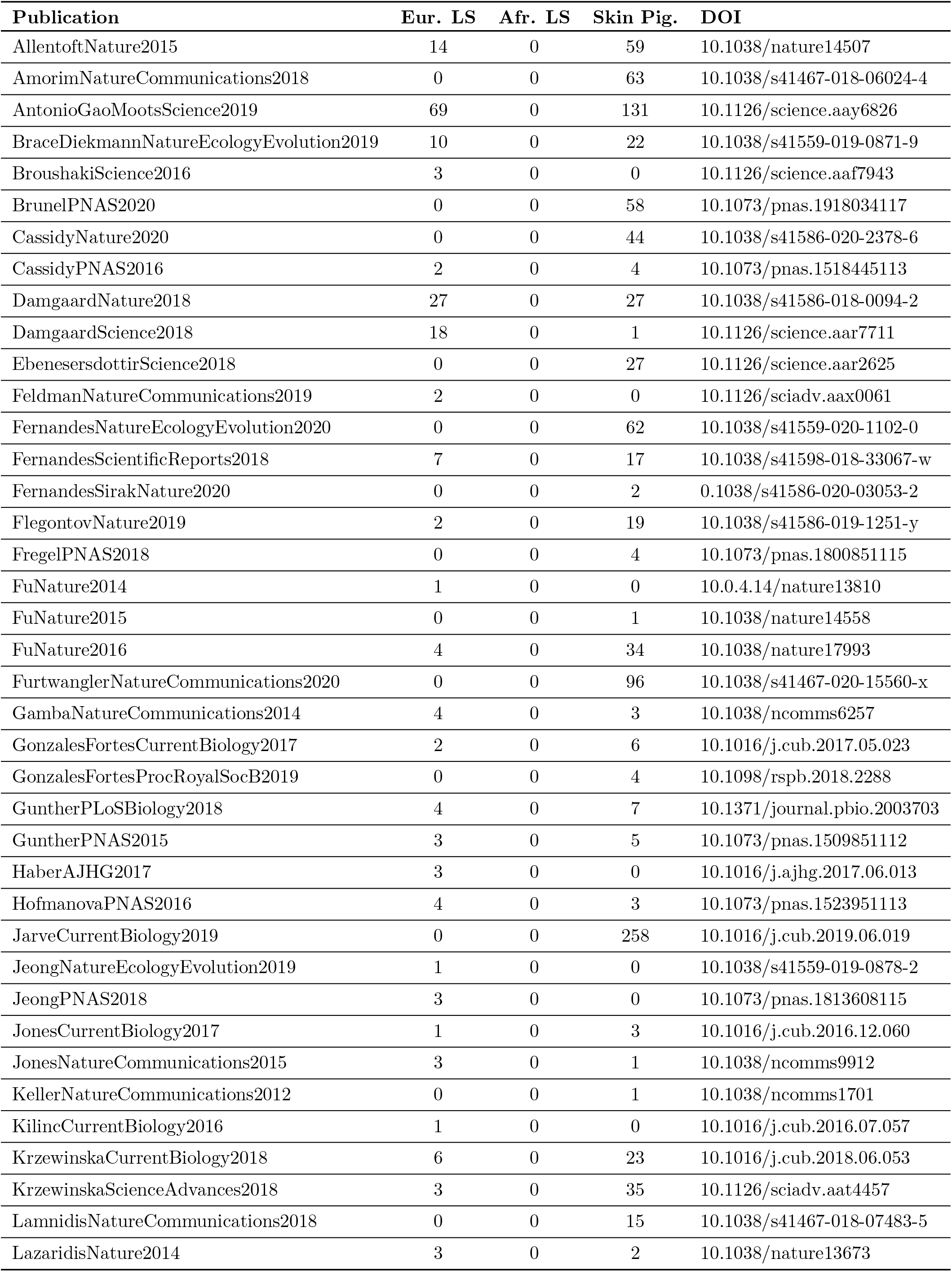
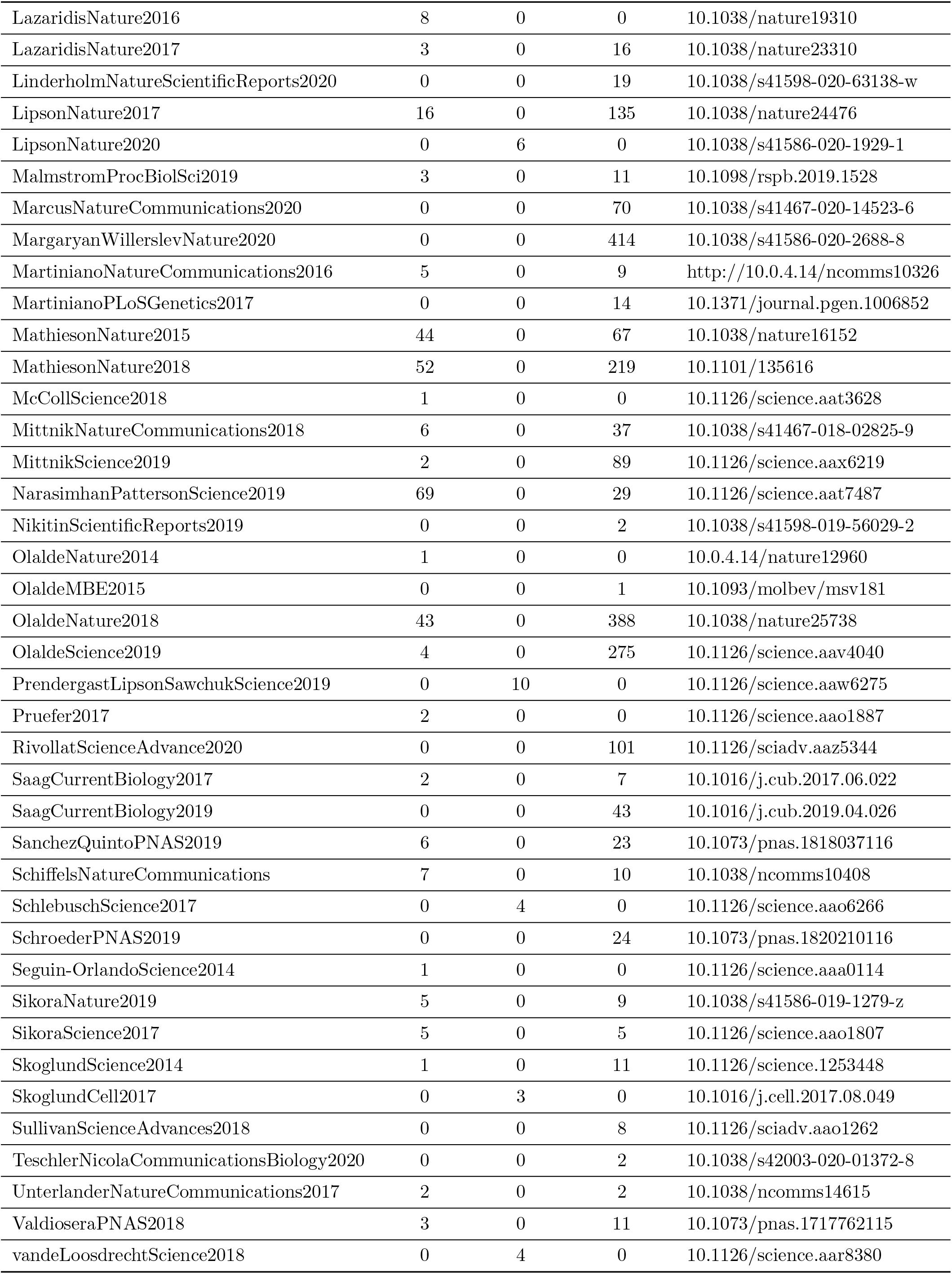
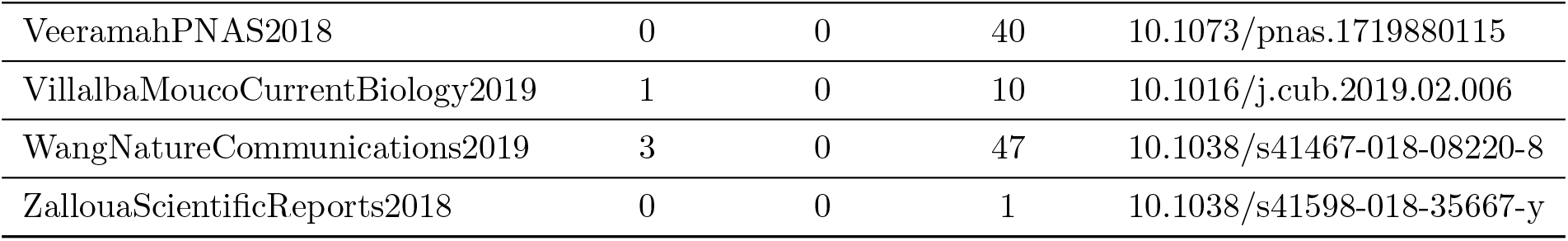
Original publications for samples used in each analysis. Publications are identified by First author name, year, and journal, with the doi provided. Also given are the number of samples from that publication used in each analysis (Eurasian Lifestyle, African *FADS1* Lifestyle, Skin Pigmentation). Specific samples used can be found in Supplemental File S1

## Bibliography

Ameur, A., Enroth, S., Johansson, A., Zaboli, G., Igl, W., Johansson, A. C. V., Rivas, M. A., Daly, M. J., Schmitz, G., Hicks, A. A., Meitinger, T., Feuk, L., van Duijn, C., Oostra, B., Pramstaller, P. P., Rudan, I., Wright, A. F., Wilson, J. F., Campbell, H., and Gyllensten, U. 2012. Genetic adaptation of fatty-acid metabolism: a human-specific haplotype increasing the biosynthesis of long-chain omega-3 and omega-6 fatty acids. American journal of human genetics, 90(5): 809–820.

Barrios-Correa, A. A., Estrada, J. A., and Contreras, I. 2018. Leptin Signaling in the Control of Metabolism and Appetite: Lessons from Animal Models. Journal of Molecular Neuroscience, 66(3): 390–402.

Benton, M. L., Abraham, A., LaBella, A. L., Abbot, P., Rokas, A., and Capra, J. A. 2021. The influence of evolutionary history on human health and disease. Nature Reviews Genetics, 22(5): 269–283.

Berg, J. J. and Coop, G. 2014. A Population Genetic Signal of Polygenic Adaptation. PLoS Genetics, 10(8).

Buckley, M. T., Racimo, F., Allentoft, M. E., Jensen, M. K., Jonsson, A., Huang, H., Hormozdiari, F., Sikora, M., Marnetto, D., Eskin, E., Jørgensen, M. E., Grarup, N., Pedersen, O., Hansen, T., Kraft, P., Willerslev, E., and Nielsen, R. 2017. Selection in Europeans on Fatty Acid Desaturases Associated with Dietary Changes. Molecular biology and evolution, 34(6): 1307–1318.

Catassi, C. and Catassi, G. N. 2018. The puzzling relationship between human leukocyte antigen HLA genes and celiac disease. Saudi journal of gastroenterology : official journal of the Saudi Gastroenterology Association, 24(5): 257–258.

Chaki, M., Sengupta, M., Mondal, M., Bhattacharya, A., Mallick, S., Bhadra, R., Consortium, I. G. V., and Ray, K. 2011. Molecular and Functional Studies of Tyrosinase Variants Among Indian Oculocutaneous Albinism Type 1 Patients. Journal of Investigative Dermatology, 131(1): 260–262.

Chen, J., Swofford, R., Johnson, J., Cummings, B. B., Rogel, N., Lindblad-Toh, K., Haerty, W., Di Palma, F., and Regev, A. 2018. A quantitative framework for characterizing the evolutionary history of mammalian gene expression. Genome Research, 29(1): 53–63.

Chou, S., Upton, H., Bao, K., Schulze-Gahmen, U., Samelson, A. J., He, N., Nowak, A., Lu, H., Krogan, N. J., Zhou, Q., and Alber, T. 2013. HIV-1 Tat recruits transcription elongation factors dispersed along a flexible AFF4 scaffold. Proceedings of the National Academy of Sciences of the United States of America, 110(2): E123–31.

Cohen, L., Henzel, W. J., and Baeuerle, P. A. 1998. IKAP is a scaffold protein of the IkappaB kinase complex. Nature, 395(6699): 292–296.

Colbran, L. L., Gamazon, E. R., Zhou, D., Evans, P., Cox, N. J., and Capra, J. A. 2019. Inferred divergent gene regulation in archaic hominins reveals potential phenotypic differences. Nature Ecology & Evolution.

Console, L., Scalise, M., Tonazzi, A., Giangregorio, N., and Indiveri, C. 2018. Characterization of Exosomal SLC22A5 (OCTN2) carnitine transporter. Scientific Reports, 8(1): 3758.

Das, S., Forer, L., Schönherr, S., Sidore, C., Locke, A. E., Kwong, A., Vrieze, S. I., Chew, E. Y., Levy, S., McGue, M., Schlessinger, D., Stambolian, D., Loh, P.-R., Iacono, W. G., Swaroop, A., Scott, L. J., Cucca, F., Kronenberg, F., Boehnke, M., Abecasis, G. R., and Fuchsberger, C. 2016. Next-generation genotype imputation service and methods. Nature genetics, 48(10): 1284–1287.

De Silvestri, A., Capittini, C., Poddighe, D., Valsecchi, C., Marseglia, G., Tagliacarne, S. C., Scotti, V., Rebuffi, C., Pasi, A., Martinetti, M., and Tinelli, C. 2018. HLA-DQ genetics in children with celiac disease: a meta-analysis suggesting a two-step genetic screening procedure starting with HLA-DQ *β* chains. Pediatric Research, 83(3): 564–572.

Dehghani, M. R., Mehrjardi, M. Y. V., Dilaver, N., Tajamolian, M., Enayati, S., Ebrahimi, P., Amoli, M. M., Farooqi, S., and Maroofian, R. 2018. Potential role of gender specific effect of leptin receptor deficiency in an extended consanguineous family with severe early-onset obesity. European Journal of Medical Genetics, 61(8): 465–467.

Devlin, B. and Roeder, K. 1999. Genomic Control for Association Studies. Biometrics, 55(4): 997–1004.

D’Mello, S. A. N., Finlay, G. J., Baguley, B. C., and Askarian-Amiri, M. E. 2016. Signaling Pathways in Melanogenesis. International journal of molecular sciences, 17(7): 1144.

Eisenberg, E. and Levanon, E. Y. 2003. Human housekeeping genes are compact. Trends in Genetics, 19(7): 362–365.

Eisenberg, E. and Levanon, E. Y. 2013. Human housekeeping genes, revisited. Trends in Genetics, 29(10): 569–574.

Enard, D., Cai, L., Gwennap, C., and Petrov, D. A. 2016. Viruses are a dominant driver of protein adaptation in mammals. eLife, 5: e12469.

Engelken, J., Espadas, G., Mancuso, F. M., Bonet, N., Scherr, A.-L., Jímenez-Álvarez, V., Codina-Solà, M., Medina-Stacey, D., Spataro, N., Stoneking, M., Calafell, F., Sabidó, E., and Bosch, E. 2016. Signatures of Evolutionary Adaptation in Quantitative Trait Loci Influencing Trace Element Homeostasis in Liver. Molecular Biology and Evolution, 33(3): 738–754.

Farooqi, I. S., Wangensteen, T., Collins, S., Kimber, W., Matarese, G., Keogh, J. M., Lank, E., Bottomley, B., Lopez-Fernandez, J., Ferraz-Amaro, I., Dattani, M. T., Ercan, O., Myhre, A. G., Retterstol, L., Stanhope, R., Edge, J. A., McKenzie, S., Lessan, N., Ghodsi, M., De Rosa, V., Perna, F., Fontana, S., Barroso, I., Undlien, D. E., and O’Rahilly, S. 2007. Clinical and molecular genetic spectrum of congenital deficiency of the leptin receptor. The New England journal of medicine, 356(3): 237–247.

Field, Y., Boyle, E. A., Telis, N., Gao, Z., Gaulton, K. J., Golan, D., Yengo, L., Rocheleau, G., Froguel, P., McCarthy, M. I., and Pritchard, J. K. 2016. Detection of human adaptation during the past 2000 years. Science (New York, N.Y.), 354(6313): 760–764.

Flanagan, J. L., Simmons, P. A., Vehige, J., Willcox, M. D. P., and Garrett, Q. 2010. Role of carnitine in disease. Nutrition & Metabolism, 7(1): 30.

Fu, Q., Hajdinjak, M., Moldovan, O. T., Constantin, S., Mallick, S., Skoglund, P., Patterson, N., Rohland, N., Lazaridis, I., Nickel, B., Viola, B., Prüfer, K., Meyer, M., Kelso, J., Reich, D., and Pääbo, S. 2015. An early modern human from Romania with a recent Neanderthal ancestor. Nature, 524(7564): 216–219.

Gamazon, E. R., Wheeler, H. E., Shah, K. P., Mozaffari, S. V., Aquino-Michaels, K., Carroll, R. J., Eyler, A. E., Denny, J. C., GTEx Consortium, Nicolae, D. L., Cox, N. J., and Im, H. K. 2015. A gene-based association method for mapping traits using reference transcriptome data. Nat Genet, 47(9): 1091–1098.

Ghodsinejad Kalahroudi, V., Kamalidehghan, B., Arasteh Kani, A., Aryani, O., Tondar, M., Ahmadipour, F., Chung, L. Y., and Houshmand, M. 2014. Two Novel Tyrosinase (TYR) Gene Mutations with Pathogenic Impact on Oculocutaneous Albinism Type 1 (OCA1). PLOS ONE, 9(9): e106656.

Goude, G. and Fontugne, M. 2016. Carbon and nitrogen isotopic variability in bone collagen during the Neolithic period: Influence of environmental factors and diet. Journal of Archaeological Science, 70: 117–131.

Grossman, S. R., Andersen, K. G., Shlyakhter, I., Tabrizi, S., Winnicki, S., Yen, A., Park, D. J., Griesemer, D., Karlsson, E. K., Wong, S. H., Cabili, M., Adegbola, R. A., Bamezai, R. N., Hill, A. V., Vannberg, F. O., Rinn, J. L., Lander, E. S., Schaffner, S. F., and Sabeti, P. C. 2013. Identifying Recent Adaptations in Large-Scale Genomic Data. Cell, 152(4): 703–713.

Guillin, O. M., Vindry, C., Ohlmann, T., and Chavatte, L. 2019. Selenium, Selenoproteins and Viral Infection.

Haak, W., Lazaridis, I., Patterson, N., Rohland, N., Mallick, S., Llamas, B., Brandt, G., Nordenfelt, S., Harney, E., Stewardson, K., Fu, Q., Mittnik, A., Bánffy, E., Economou, C., Francken, M., Friederich, S., Pena, R. G., Hallgren, F., Khartanovich, V., Khokhlov, A., Kunst, M., Kuznetsov, P., Meller, H., Mochalov, O., Moiseyev, V., Nicklisch, N., Pichler, S. L., Risch, R., Rojo Guerra, M. A., Roth, C., Szécsényi-Nagy, A., Wahl, J., Meyer, M., Krause, J., Brown, D., Anthony, D., Cooper, A., Alt, K. W., and Reich, D. 2015. Massive migration from the steppe was a source for Indo-European languages in Europe. Nature, 522(7555): 207–211.

Hancock, A. M., Witonsky, D. B., Gordon, A. S., Eshel, G., Pritchard, J. K., Coop, G., and Di Rienzo, A. 2008. Adaptations to climate in candidate genes for common metabolic disorders. PLoS genetics, 4(2): e32–e32.

He, N., Liu, M., Hsu, J., Xue, Y., Chou, S., Burlingame, A., Krogan, N. J., Alber, T., and Zhou, Q. 2010. HIV-1 Tat and host AFF4 recruit two transcription elongation factors into a bifunctional complex for coordinated activation of HIV-1 transcription. Molecular cell, 38(3): 428–438.

Irving-Pease, E. K., Muktupavela, R., Dannemann, M., and Racimo, F. 2021. Quantitative paleogenetics: what can ancient dna tell us about complex trait evolution? arXiv.

Ju, D. and Mathieson, I. 2020. The evolution of skin pigmentation associated variation in West Eurasia. bioRxiv, page 2020.05.08.085274.

Kentish, S. J., Wittert, G. A., Blackshaw, L. A., and Page, A. J. 2013. A chronic high fat diet alters the homologous and heterologous control of appetite regulating peptide receptor expression. Peptides, 46: 150–158.

Lamason, R. L., Mohideen, M.-A. P. K., Mest, J. R., Wong, A. C., Norton, H. L., Aros, M. C., Jurynec, M. J., Mao, X., Humphreville, V. R., Humbert, J. E., Sinha, S., Moore, J. L., Jagadeeswaran, P., Zhao, W., Ning, G., Makalowska, I., McKeigue, P. M., O’donnell, D., Kittles, R., Parra, E. J., Mangini, N. J., Grunwald, D. J., Shriver, M. D., Canfield, V. A., and Cheng, K. C. 2005. SLC24A5, a putative cation exchanger, affects pigmentation in zebrafish and humans. Science (New York, N.Y.), 310(5755): 1782–1786.

Lek, M., Karczewski, K. J., Minikel, E. V., Samocha, K. E., Banks, E., Fennell, T., O’Donnell-Luria, A. H., Ware, J. S., Hill, A. J., Cummings, B. B., Tukiainen, T., Birnbaum, D. P., Kosmicki, J. A., Duncan, L. E., Estrada, K., Zhao, F., Zou, J., Pierce-Hoffman, E., Berghout, J., Cooper, D. N., Deflaux, N., DePristo, M., Do, R., Flannick, J., Fromer, M., Gauthier, L., Goldstein, J., Gupta, N., Howrigan, D., Kiezun, A., Kurki, M. I., Moonshine, A. L., Natarajan, P., Orozco, L., Peloso, G. M., Poplin, R., Rivas, M. A., Ruano-Rubio, V., Rose, S. A., Ruderfer, D. M., Shakir, K., Stenson, P. D., Stevens, C., Thomas, B. P., Tiao, G., Tusie-Luna, M. T., Weisburd, B., Won, H.-H., Yu, D., Altshuler, D. M., Ardissino, D., Boehnke, M., Danesh, J., Donnelly, S., Elosua, R., Florez, J. C., Gabriel, S. B., Getz, G., Glatt, S. J., Hultman, C. M., Kathiresan, S., Laakso, M., McCarroll, S., McCarthy, M. I., McGovern, D., McPherson, R., Neale, B. M., Palotie, A., Purcell, S. M., Saleheen, D., Scharf, J. M., Sklar, P., Sullivan, P. F., Tuomilehto, J., Tsuang, M. T., Watkins, H. C., Wilson, J. G., Daly, M. J., MacArthur, D. G., and Consortium, E. A. 2016. Analysis of protein-coding genetic variation in 60,706 humans. Nature, 536: 285.

Li, R., Chen, Y., Ritchie, M. D., and Moore, J. H. 2020. Electronic health records and polygenic risk scores for predicting disease risk. Nature Reviews Genetics, 21(8): 493–502.

Liao, Y., Wang, J., Jaehnig, E. J., Shi, Z., and Zhang, B. 2019. WebGestalt 2019: gene set analysis toolkit with revamped UIs and APIs. Nucleic acids research, 47(W1): W199–W205.

Loos, R. J. F., Rankinen, T., Chagnon, Y., Tremblay, A., Pérusse, L., and Bouchard, C. 2006. Polymorphisms in the leptin and leptin receptor genes in relation to resting metabolic rate and respiratory quotient in the Québec Family Study. International Journal of Obesity, 30(1): 183–190.

Luca, F., Perry, G. H., and Di Rienzo, A. 2010. Evolutionary adaptations to dietary changes. Annual review of nutrition, 30: 291–314.

Machiela, M. J. and Chanock, S. J. 2015. LDlink: a web-based application for exploring population-specific haplotype structure and linking correlated alleles of possible functional variants. Bioinformatics (Oxford, England), 31(21): 3555–3557.

Mancini, A., Koch, A., Whetton, A. D., and Tamura, T. 2004. The M-CSF receptor substrate and interacting protein FMIP is governed in its subcellular localization by protein kinase C-mediated phosphorylation, and thereby potentiates M-CSF-mediated differentiation. Oncogene, 23(39): 6581–6589.

Mann, N. J. 2018. A brief history of meat in the human diet and current health implications. Meat Science, 144: 169–179.

Marciniak, S. and Perry, G. H. 2017. Harnessing ancient genomes to study the history of human adaptation. Nature Reviews Genetics, 18(11): 659–674.

Mathieson, S. and Mathieson, I. 2018. FADS1 and the timing of human adaptation to agriculture. Molecular Biology and Evolution, 35(12): 2957–2970.

Norman, C. S., O’Gorman, L., Gibson, J., Pengelly, R. J., Baralle, D., Ratnayaka, J. A., Griffiths, H., Rose-Zerilli, M., Ranger, M., Bunyan, D., Lee, H., Page, R., Newall, T., Shawkat, F., Mattocks, C., Ward, D., Ennis, S., and Self, J. E. 2017. Identification of a functionally significant tri-allelic genotype in the Tyrosinase gene (TYR) causing hypomorphic oculocutaneous albinism (OCA1B). Scientific Reports, 7(1): 4415.

Okoro, P. C., Schubert, R., Guo, X., Johnson, W. C., Rotter, J. I., Hoeschele, I., Liu, Y., Im, H. K., Luke, A., Dugas, L. R., and Wheeler, H. E. 2021. Transcriptome prediction performance across machine learning models and diverse ancestries. Human Genetics and Genomics Advances.

Olsson, O. and Paik, C. 2016. Long-run cultural divergence: Evidence from the Neolithic Revolution. Journal of Development Economics, 122: 197–213.

Petty, L. E., Highland, H. M., Gamazon, E. R., Hu, H., Karhade, M., Chen, H.-H., de Vries, P. S., Grove, M. L., Aguilar, D., Bell, G. I., Huff, C. D., Hanis, C. L., Doddapaneni, H., Munzy, D. M., Gibbs, R. A., Ma, J., Parra, E. J., Cruz, M., Valladares-Salgado, A., Arking, D. E., Barbeira, A., Im, H. K., Morrison, A. C., Boerwinkle, E., and Below, J. E. 2019. Functionally oriented analysis of cardiometabolic traits in a trans-ethnic sample. Human Molecular Genetics, 00(00): 1–13.

Pierini, F. and Lenz, T. L. 2018. Divergent Allele Advantage at Human MHC Genes: Signatures of Past and Ongoing Selection. Molecular Biology and Evolution, 35(9): 2145–2158.

Rees, J. S., Castellano, S., and Andrés, A. M. 2020. The Genomics of Human Local Adaptation. Trends in Genetics, 36(6): 415–428.

Skoglund, P. and Mathieson, I. 2018. Ancient Human Genomics: The First Decade. Annu. Rev. Genom. Hum. Genet, 198(April): 1–824.

Soejima, M. and Koda, Y. 2007. Population differences of two coding SNPs in pigmentation-related genes SLC24A5 and SLC45A2. International journal of legal medicine, 121(1): 36–39.

Sturm, R. A. and Duffy, D. L. 2012. Human pigmentation genes under environmental selection. Genome Biology, 13(9): 248.

Tamura, T., Mancini, A., Joos, H., Koch, A., Hakim, C., Dumanski, J., Weidner, K. M., and Niemann, H. 1999. FMIP, a novel Fms-interacting protein, affects granulocyte/macrophage differentiation. Oncogene, 18(47): 6488–6495.

The 1000 Genomes Project Consortium 2015. A global reference for human genetic variation. Nature, 526(7571): 68–74.

Voight, B. F., Kudaravalli, S., Wen, X., and Pritchard, J. K. 2006. A Map of Recent Positive Selection in the Human Genome. PLOS Biology, 4(3): e72.

Weiss, C. V., Harshman, L., Inoue, F., Fraser, H. B., Petrov, D. A., Ahituv, N., and Gokhman, D. 2021. The cis-regulatory effects of modern human-specific variants. Elife, 10.

White, L., Romagné, F., Müller, E., Erlebach, E., Weihmann, A., Parra, G., Andrés, A. M., and Castellano, S. 2015. Genetic Adaptation to Levels of Dietary Selenium in Recent Human History. Molecular Biology and Evolution, 32(6): 1507–1518.

Wilde, S., Timpson, A., Kirsanow, K., Kaiser, E., Kayser, M., Unterländer, M., Hollfelder, N., Potekhina, D., Schier, W., Thomas, M. G., and Burger, J. 2014. Direct evidence for positive selection of skin, hair, and eye pigmentation in Europeans during the last 5,000 y. Proceedings of the National Academy of Sciences, page 201316513.

Ye, K., Gao, F., Wang, D., Bar-Yosef, O., and Keinan, A. 2017. Dietary adaptation of FADS genes in Europe varied across time and geography. Nature Ecology & Evolution, 1(7): 167.

Zhang, T., Choi, J., Kovacs, M., Shi, J., Xu, M., Goldstein, A., Iles, M., Duffy, D., MacGregor, S., Amundadottir, L., Law, M., Loftus, S., Pavan, W., and Brown, K. 2017. Cell-type specific eQTL of primary melanocytes facilitates identification of melanoma susceptibility genes. Cell-type-specific eQTL of primary melanocytes facilitates identification of melanoma susceptibility genes, page 231423.

Zheng, W.-S., He, Y.-X., Cui, C.-Y., Ouzhu, L., Deji, Q., Peng, Y., Bai, C.-J., Duoji, Z., Gongga, L., Bian, B., Baima, K., Pan, Y.-Y., Qu, L., Kang, M., Ciren, Y., Baima, Y., Guo, W., Yang, L., Zhang, H., Zhang, X.-M., Guo, Y.-B., Xu, S.-H., Chen, H., Zhao, S.-G., Cai, Y., Liu, S.-M., Wu, T.-Y., Qi, X.-B., and Su, B. 2017. EP300 contributes to high-altitude adaptation in Tibetans by regulating nitric oxide production. Zoological research, 38(3): 163–170.

Zhou, D., Jiang, Y., Zhong, X., Cox, N. J., Liu, C., and Gamazon, E. R. 2020. A unified framework for joint-tissue transcriptome-wide association and Mendelian randomization analysis. Nature genetics.

Zhu, H. and Zhou, X. 2020. Transcriptome-wide association studies: a view from Mendelian randomization. Quantitative Biology.

